# Causal inference in environmental sound recognition

**DOI:** 10.1101/2020.07.13.200949

**Authors:** James Traer, Sam V. Norman-Haignere, Josh H. McDermott

## Abstract

Sound is caused by physical events in the world. Do humans infer these causes when recognizing sound sources? We tested whether the recognition of common environmental sounds depends on the inference of a basic physical variable – the source intensity (i.e., the power that produces a sound). A source’s intensity can be inferred from the intensity it produces at the ear and its distance, which is normally conveyed by reverberation. Listeners could thus use intensity at the ear and reverberation to constrain recognition by inferring the underlying source intensity. Alternatively, listeners might separate these acoustic cues from their representation of a sound’s identity in the interest of invariant recognition. We compared these two hypotheses by measuring recognition accuracy for sounds with typically low or high source intensity (e.g., pepper grinders vs. trucks) that were presented across a range of intensities at the ear or with reverberation cues to distance. The recognition of low-intensity sources (e.g., pepper grinders) was impaired by high presentation intensities or reverberation that conveyed distance, either of which imply high source intensity. Neither effect occurred for high-intensity sources. The results suggest that listeners implicitly use the intensity at the ear along with distance cues to infer a source’s power and constrain its identity. The recognition of real-world sounds thus appears to depend upon the inference of their physical generative parameters, even generative parameters whose cues might otherwise be separated from the representation of a sound’s identity.

## 1. Introduction

Just by listening, we can tell that we are walking next to a stream, that a mosquito is hovering nearby, or that an animal is growling. Though it is clear that humans can recognize environmental sounds (Balas, 1993; Giordano, 2003; Gygi, Kidd, & Watson, 2004, 2007; Gygi & Shafiro, 2011; Leech, Gygi, Aydelott, & Dick, 2009; Lemaitre & Heller, 2013; McDermott & Simoncelli, 2011), the underlying computations remain poorly understood.

A central challenge of recognition is that similar entities in the world can produce very different sensory signals, as when an object is viewed under different lighting conditions, or a sound is heard from near or far (Fig. 1A). Somehow our sensory systems must generalize across this variation while retaining the ability to discriminate different objects (Carruthers et al., 2015; DiCarlo & Cox, 2007; Liu, Montes-Lourido, Wang, & Sadagopan, 2019; Rust & DiCarlo, 2010; Sharpee, Atencio, & Schreiner, 2011). One possibility is that listeners separate or remove unwanted variation from a sound’s internal representation to achieve invariance, akin to how contemporary machine recognition systems are believed to associate sets of stimuli with labels (Goodfellow, Lee, Le, Saxe, & Ng, 2009; Tacchetti, Isik, & Poggio, 2018). In speech, variation in word acoustics due to speaking speed as well as the pitch and vocal tract of the speaker (Allen, Miller, & DeSteno, 2003; Hillenbrand, Getty, Clark, & Wheeler, 1995; Stevens, 2000) is often thought to be normalized or separated from the representation of speech content (Holt, 2006; Johnson, 2005; Lehet & Holt, 2020; Nusbaum & Magnuson, 1997; Pisoni, 1997). Similar normalization mechanisms could underlie representations of melodies, the recognition of which is also robust to time dilation, pitch transposition, and other transformations (Attneave & Olson, 1971; Dowling & Fujitani, 1970; McDermott, Lehr, & Oxenham, 2008). Background noise (Ding & Simon, 2013; Kell & McDermott, 2019; Khalighinejad, Herrero, Mehta, & Mesgarani, 2019; Moore, Lee, & Theunissen, 2013; Rabinowitz, Willmore, King, & Schnupp, 2013; Scott & McGettigan, 2013), reverberation (Mesgarani, David, Fritz, & Shamma, 2014; Traer & McDermott, 2016) and intensity (Billimoria, Kraus, Narayan, Maddox, & Sen, 2008; Sadagopan & Wang, 2008) may also be partially separated from the representation of a sound’s source.

**Figure 1.**
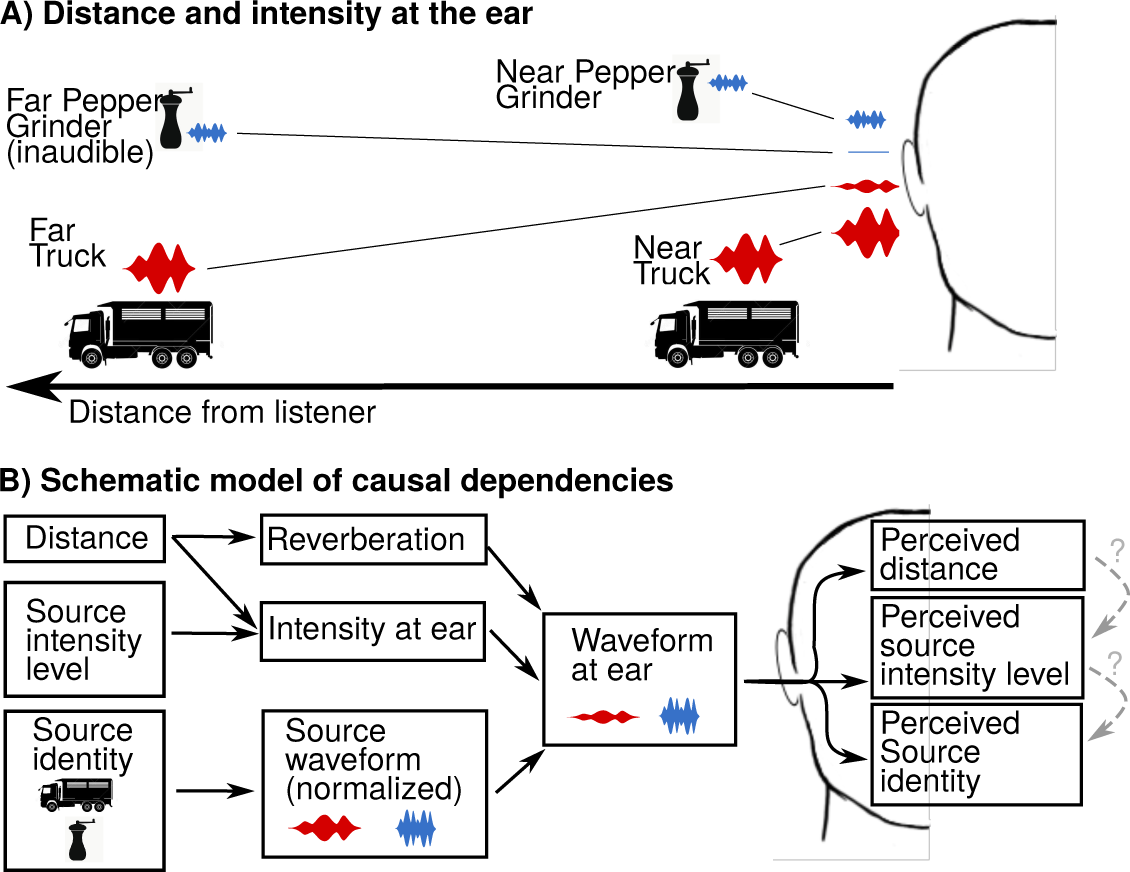
Overview and hypotheses. (A) Schematic depicting the impact of source intensity and distance on the intensity at the ear: while sounds with high-intensity sources (e.g., a truck) induce a wide range of ear intensities depending on the distance (due to the inverse-square law), low-intensity sources (e.g., pepper grinder) never produce high ear intensities. (B) A graphical model of the interdependencies of the acoustic features we investigate: solid-lines represent causal relationships in the generative process, dashed-lines represent hypothetical relationships humans may use to infer the sound source. One hypothesis is that source identity, intensity and distance are each inferred separately. Another possibility is that that these judgments inform each other. We use manipulations of sound intensity and reverberation to investigate whether source recognition is affected by perceived source intensity, which might be inferred from perceived distance.

However, invariant association of labels with stimuli is not the only goal of perception. In the case of audition, most everyday sounds are caused by physical interactions (e.g., impacts, scrapes, fracturing, fluid motion etc.) (Gaver, 1993). Listeners have some ability to describe these interactions for common sound sources (Conan et al., 2014; Grassi, 2005; Grassi, Pastore, & Lemaitre, 2013; Guyot et al., 2017; Hjortkjær & McAdams, 2016; Lemaitre & Heller, 2012; Lutfi, 2008; Rocchesso & Fontana, 2003; Traer, Cusimano, & McDermott, 2019), raising the possibility that the inference of the mechanisms that compose the source might contribute importantly to everyday recognition even if they are not explicitly part of the verbal label with which a sound is identified.

To investigate these possibilities, we examined the effect of intensity on sound recognition as a simple test case (Fig. 1B). A sound’s intensity at the ear depends upon the both the source’s intensity (which depends on the nature of the source) and the distance to the listener, due to the inverse-square law (Fig 1B). These relationships are occasionally violated in modern listening conditions due to electrical amplification of sound, but are nonetheless present for most of our daily auditory experience. It is clear from everyday experience that listeners can recognize sound sources, and that they have some ability to judge a source’s intensity, and to estimate source distance. However, the relationship between the processes underlying these judgments is poorly understood. They could be largely distinct, with recognition that is invariant to intensity and distance cues. Alternatively, humans might jointly infer sources, their intensities, and their arrangement within the scene. The latter hypothesis predicts that judgments of one parameter should tacitly affect another. For example, inferred distance could affect judgments of source intensity, which in turn could affect source recognition (Fig. 1C). Some evidence that intensity might influence recognition comes from the finding that listeners are biased by intensity in sound memory tasks (Susini, Houix, Seropian, & Lemaitre, 2019), but to our knowledge the effect on recognition of everyday sounds had not been explicitly measured prior to this study.

In Experiments 1—9 we investigated how source recognition was affected by sound intensity at the ear, and by reverberation, which conveys source distance and could thus indirectly influence the inferred intensity of a source. We compared the effects for high- and low-intensity sources (e.g., a truck and a pepper grinder). Our results show that listeners consistently misidentify low-intensity sources when presented with either high intensities at the ear or reverberation conveying distance, both of which entail an implausibly high source intensity. This result contradicts the hypothesis that recognition is invariant to intensity and reverberation, but is consistent with causal inference, because neither high-intensity nor distant sounds can possibly be generated by low-intensity sources. Experiments 1-6 were run with a large set of sound recordings made in natural scenes. To ensure that our results were robust to the reverberation intrinsic to these natural recordings, in Experiments 7-9 we replicated key results with a set of studio-recordings that had minimal reverberation.

To address the possibility that the result could instead be driven by unfamiliar combinations of acoustic cues, with low-intensity sources misidentified when they are encountered in conditions that have plausibly not been previously encountered by participants (i.e., at high intensities or with reverberation appropriate for enclosed spaces), we conducted two follow-up experiments. In Experiments 10 and 11 we explicitly tested whether the reverberation we applied sounded appropriate for our recorded sounds. We found that the reverberation was heard to be less appropriate for low-intensity sound sources typically encountered outdoors than those typically encountered indoors, consistent with the lower reverberation found in outdoor environments (Traer & McDermott, 2016). However, when the source recognition experiments (Experiments 4 and 8) were reanalyzed separately for indoor and outdoor sounds, low-intensity outdoor sounds were no more frequently misidentified than indoor sounds. Humans thus misidentify sources under conditions that are physically impossible in natural conditions, but not conditions that are acoustically atypical, a result that provides additional support for causal inference.

Finally, we replicated the key results with experimental variants designed to rule out various potential confounds. We show that the main results cannot be explained by intrinsic differences in our sound sources, such as spectral content, or the artificial amplification of typically inaudible structure (Experiment 12).

These results suggest that an interwoven set of inferences underlie everyday recognition: listeners infer distance from reverberation, judge source intensity from the inferred distance and intensity at the ear, and then identify sources in part based on the inferred source intensity (Fig. 1C). Sound recognition thus appears to be intrinsically linked to intuitive causal inference of the scene and source properties.

## 2. Experiments 1 and 2: Everyday Sound Recognition is Not Invariant to Intensity

We began by measuring the ability of listeners to recognize everyday sounds presented to the ear at different intensities. We then assessed whether the typical intensity of the sound source in the real world affected listeners’ performance.

To assess identification accuracy, we first used an “open set” recognition task (Experiment 1): on each trial listeners heard a 2-second sound and were asked to type a description of it (e.g., “Hand grinder. For spices or coffee beans.”; Fig. 2A, top). We adopted this methodology because it is more ecologically relevant than a forced-choice task in which listeners are presented with a fixed set of options (as have been used in many previous studies of environmental sound recognition (Gygi et al., 2004; Gygi & Shafiro, 2011; McDermott & Simoncelli, 2011)). We were also concerned that affording listeners a set of possible sound identities might artificially boost performance and mask differences between conditions that might otherwise be present in real-world conditions. To assess the accuracy of the descriptions provided by each listener, we had online workers guess the sound heard by each listener based on these descriptions (via Amazon’s Mechanical Turk platform; Fig. 2A, bottom). The online workers, who did not hear the sound, chose the sound label that best fit the listener’s description from a list of 10 possible choices. The accuracy of a listener’s descriptions was quantified as the fraction of trials on which the online workers were able to correctly identify the sound from their descriptions.

**Figure 2.**
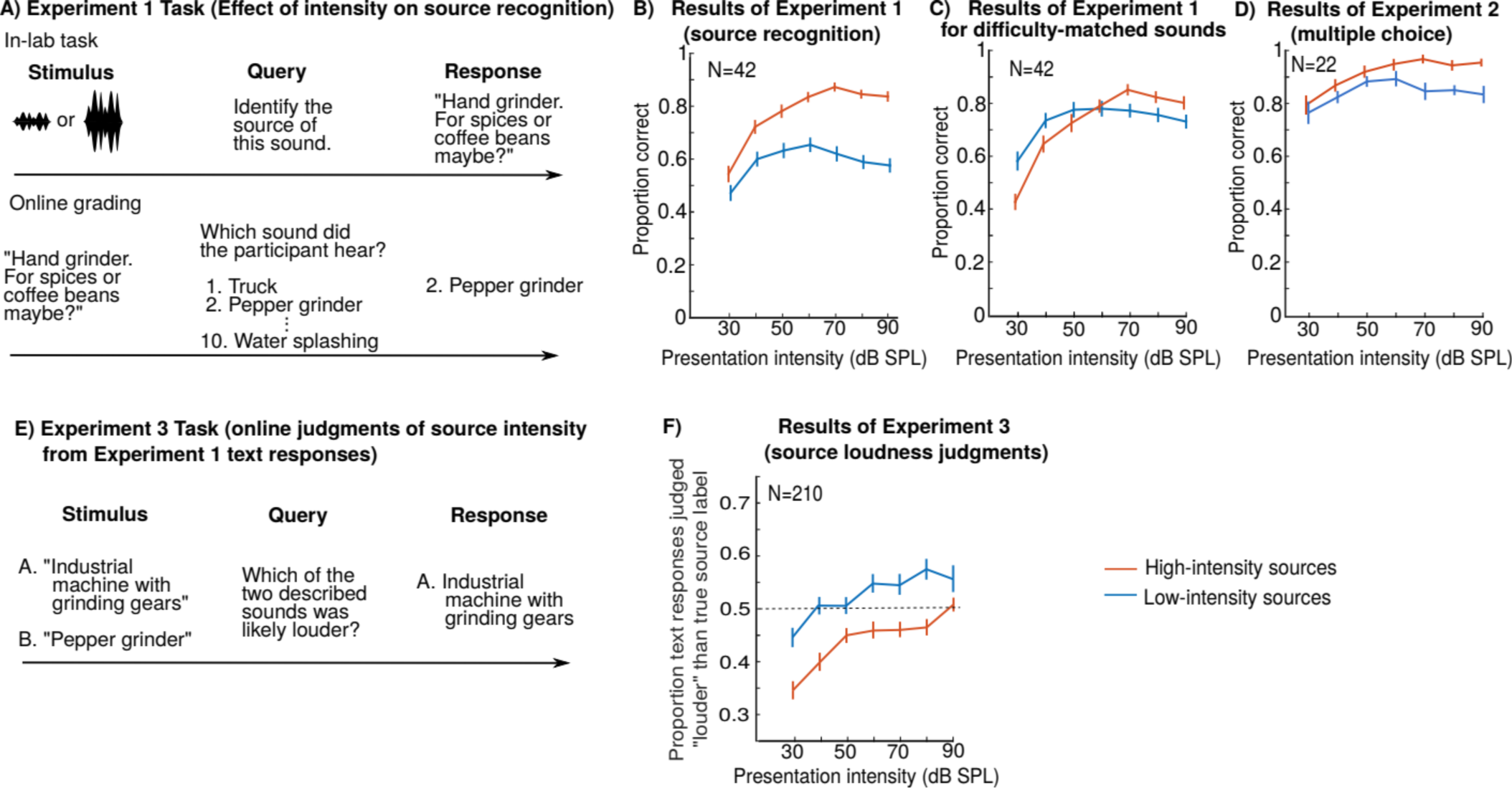
Experiments 1-3: Misidentification of sound sources presented with implausible intensities or reverberation suggests casual inference. (A) Schematic of the open-set recognition task used in Experiment 1, which was intended to be more sensitive and ecologically valid than a traditional forced-choice experiment. In-lab participants (top) heard sounds at different presentation intensities and typed descriptions of what they heard, here with the sound of a pepper grinder as an example. A separate group of online workers (bottom) tried to identify the sound the in-lab participants heard from a list of 10 possible choices using only the written descriptions of the in-lab participants. (B) Results of Experiment 1: Recognition accuracy for low-intensity and high-intensity sources as a function of the physical intensity at which they were presented to the ear. Recognition accuracy reflects the fraction of online graders who correctly guessed the sound from the in-lab descriptions. Error bars show one standard error of the mean across participants from the in-lab experiment. Recognition declined for low-intensity sources when presented above 60dB SPL, which is inconsistent with the invariance hypothesis. (C) Results of Experiment 1 with overall recognition accuracy equated for low-intensity and high-intensity sources. Difficulty was equated by removing the easiest-to-recognize high-intensity sources and the hardest-to-recognize low-intensity sources (averaging across presentation intensity). The interaction between source intensity and presentation intensity persisted when controlling for overall difficulty. (D) Results of Experiment 2: multiple-choice source recognition. There was again a significant interaction between source intensity and presentation intensity, but performance was higher in all conditions, as expected. (E) Schematic of the task from Experiment 3, in which online workers compared the responses of participants from Experiment 1 to the original sound label and judged which of the two corresponded to a higher intensity sound. (F) Results of Experiment 3: The fraction of typed responses from Experiment 1 that were judged to correspond to louder sources than the original label. Results are plotted for low- and high-intensity sources as a function of the presentation intensity. Dashed line represents chance performance (i.e., the point where the typed descriptions was on average judged to be no louder or quieter than the correct original sound label.

To ensure that the results of Experiment 1 were not specific to the open set recognition task, in Experiment 2 we ran the same sound recognition task with a multiple-choice (10 choices), rather than “open set”, methodology.

### 2.1. Method

All experiments were approved by the Committee on the Use of Humans as Experimental Subjects (COUHES) at MIT, and all participants gave informed consent. No participant, in-lab or online, took part in multiple experiments, ensuring that all participants were naïve to the stimuli. In-lab experiments were conducted in soundproof booths with calibrated headphones (Appendix B).

#### 2.1.1. Participants

For clarity, all participants who listened to sounds and tried to identify the source are referred to as ‘listeners’, to distinguish them from the ‘online workers’ who graded results. 42 in-lab listeners participated in Experiment 1 (22 female, 19 male; 1 listener’s gender was not recorded; mean age = 35.7 years; SD = 13.9 years). The responses from the 42 listeners from Experiment 1 were scored by 500 online workers. 22 in-lab listeners (12 female, 10 male; mean age = 25.4 years; SD = 8.8 years) participated in Experiment 2. Participants had their hearing sensitivity assessed to ensure they could adequately hear the stimuli (Appendix C). A power analysis assuming an interaction effect size of 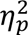=0.2 (which is on the lower end of a “large” effect size as defined by Cohen (Cohen, 1988)) indicated that 42 listeners would be needed to detect an effect of this size 80% of the time using a significance threshold of .05. The design (in which each participant heard a sound once, at one of 7 possible presentation intensities) necessitated a sample size that was a multiple of 7. We accordingly ran 42 listeners in Experiment 1, and about half as many listeners in Experiment 2 (which served as methodological control).

#### 2.1.2. Materials and procedure

Listeners were asked to identify 300 unique sounds (Table S1). The sounds were sourced from sound effects CDs and the internet, and were selected to be relatively clean and recognizable (Norman-Haignere, Kanwisher, & McDermott, 2015). The sound set included a broad range of natural sounds heard in daily life (Table S1). The sound set included some music (34 sounds) and speech stimuli (12 sounds), which might involve recognition mechanisms distinct from those for other environmental sounds (Leaver & Rauschecker, 2010; Norman-Haignere et al., 2015), but the exclusion or inclusion of these sounds did not qualitatively affect the results of the experiment. All of the sounds were 2 seconds in duration and were resampled to 20 kHz with 16-bit resolution (these were the lowest sampling rates and bit depths across the set of recordings we assembled, and so we matched all stimuli to these values). Linear ramps (10 ms) were applied to the beginning and end of each sound. All experiments manipulating intensity (Experiments 1-3 and 12) used all 300 sounds.

**Table 1.** Some example source labels and listener text responses for Experiment 1, from trials in which low-intensity sounds presented at 90dbSPL were misidentified (proportion correctly identified = 0). The 22 examples here are the trials encountered by the first 10 participants (of 42) who took part in Experiment 1.

| True source label | In-booth listener text response (Exp1) |
| --- | --- |
| velcro | rocks |
| typing | motor sound |
| flag | thunder |
| microwave | drying clothes |
| grating food | machine |
| dial tone | horn |
| chair rolling | pinball |
| slicing bread | fixing a motorcycle |
| drawer opening | pully |
| key opening door | bolt action |
| match lighting | automatic inflation or deflation |
| coloring | scribbling on chalkboard |
| soda pouring into a cup | When you have a nearly empty fountain drink with ice – those last couple of sips |
| wing flapping | Shaking something up |
| chopping food | Something being stuck/ applied to something else |
| microwave | airplane |
| leather coat | motorbike noise or Velcro ripping |
| peeling | casino machine |
| opening a letter | person noisily going-through a drawer |
| phone vibrating | horn or fire alarm |
| slicing bread | car noises –exhaust or engine |
| spray can spraying | machine noise –possibly welding tool |

Sounds were presented over headphones ranging from low (30 dB SPL) to high (90 dB SPL) in 10dB increments. Each listener heard each sound once. Across listeners, each sound was presented an equal number of times at each of the seven different intensities. Each intensity condition was presented the same number of times for each listener.

In Experiment 1, listeners were asked to type their best guess of the sound’s identity (as a single- or multi-word description), giving as much detail as possible. Because listeners were asked to identify the sound, they generally gave semantically meaningful descriptions (e.g., “clock”) rather than acoustic descriptions (e.g., “tic tic tic”). Trials were completed in four blocks of 75, between which listeners were encouraged to take a break.

To score the responses, we had online workers read descriptions from the in-lab listeners from Experiment 1. The workers did not know the task condition nor could they hear the sounds. For each description (e.g., “wind instrument playing a melody”), they were asked to identify the sound being described from a list of 10 choices drawn from the labels of the sound set used for the experiment (e.g., “Clarinet”, “Seagull”, “Drum roll”, “Violin”, etc.). The 9 foils were drawn randomly from the other sounds in the experiment.

Most of the Experiment 1 descriptions (97%) were scored by two workers. A small number were scored by 1 or 3 workers (due to a mixture of unanswered questions from the workers and idiosyncrasies in the way new questions are posted using the Mechanical Turk batch interface). Although each in-lab description was only scored by approximately 2 online workers, our analysis was based on the average recognition accuracy across listeners and across a large collection of low-intensity and high-intensity sound sources. Since there were 75 sounds for the low-intensity and high-intensity source groups, and since each sound was described by 6 in-lab listeners for each level tested, the average performance at a particular presentation intensity for either low-intensity or high-intensity sources was based on data from 450 in-lab descriptions and approximately 900 worker scorings. As a consequence, the pattern of mean recognition performance across conditions (presentation intensity x source group) was stable across independent sets of Mechanical Turk ratings (split-half Pearson correlation was 0.98).

To assess the typical source intensity for a sound, we had a different group of online workers rate how “quiet” or “loud” sounds typically are in the world on a scale of 1 (most quiet) to 10 (most loud) (Appendix A). The 25% of sounds with the lowest and highest ratings were used as the low-intensity and high-intensity sources, respectively, in the subsequent analysis (results for all quartiles are shown in Fig. S1).

#### 2.1.3 Statistics

In all experiments, repeated measures analyses of variance (ANOVAs) were used to test for main effects and interactions. The ANOVAs were performed on the proportion of trials that each listener got correct for each sound group and source condition. Mauchly’s test was used to test for violations of sphericity, and was never significant, indicating that the assumptions of the ANOVA were not violated. T-tests were used to directly compare two conditions-of-interest. In Experiment 2, in which participants scored above 75% in all conditions, the data were arc-sine transformed prior to performing statistical tests.

### 2.2 Results and discussion

If sound recognition is invariant to sound intensity, listener responses should be little affected by intensity, perhaps improving with presentation intensity due to better audibility. But if recognition instead depends upon the intensity at which sounds are encountered in natural scenes, results should differ for low-intensity and high-intensity sources. For high-intensity sources, such as a jackhammer or a lion’s roar, recognition largely increased with the presentation intensity (Fig. 2B; though there was a non-significant trend for poorer performance at the highest levels; t(41) = 1.88, p = 0.07 for comparing 70 and 90 dB). But for low-intensity sources, recognition peaked at moderate presentation intensities and then declined (t(41) = 2.85, p < 0.01 for comparing 60 and 90 dB) (intermediate trends were evident for sounds rated as having intermediate source intensities; Fig. S1). This difference produced an interaction between the effect of presentation intensity and the source intensity (F(6, 246) = 6.50, p < 0.001, 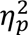=0.137, comparing low-intensity and high-intensity sources).

In the source recognition experiments (Experiment 1, Fig 2B), the low-intensity sources were less recognizable overall than the high-intensity sources (F(1, 41) = 294, p < 0.001, 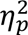=0.878). To ensure that this difference in overall recognizability could not somehow explain the interaction between presentation intensity and the source type, we equated overall recognition rates by eliminating the most-recognized and least-recognized sources in each group, respectively (we removed 25 sounds from each group, leaving 50 sounds per group in total). As is evident in Fig. 2C, the interaction between the effect of presentation intensity and the source intensity persisted after this manipulation (F(6, 246) = 7.84; p < 0.001, 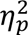=0.160).

When a multiple-choice task was used to measure recognition (Experiment 2; Fig. 2D), overall performance was higher, but the interaction between presentation intensity and real-world intensity remained (after arc-sin transforming the data: F(6, 126) = 2.29, p = 0.039). However, we note that the effect was weaker with the multiple-choice paradigm (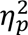=0.098), confirming our initial worry that such paradigms would artificially inflate recognition performance.

## 3. Experiment 3: Sound Descriptions are Consistent with Inferences About Source Intensity

If listeners are using intensity (either the physical intensity at the ear, or the inferred source intensity) as a cue to recognition, then when listeners misidentify low-intensity sources presented at high intensities, their erroneous answer should be a high-intensity source. To test this prediction, we analyzed the descriptive responses given for each sound in Experiment 1 (Table 1). We presented a different set of online workers with pairs of sound descriptions: a typed response from a trial in Experiment 1 along with the corresponding original source label. The online workers were asked to select the description that described a “louder” sound (Fig. 2E). We then measured the proportion of trials for which the participant’s description was judged as louder than the presented sound source, as a function of whether the source was high- or low-intensity, and the presentation intensity. If the participants tended to give descriptions of sources with about the same source intensity as the heard source, the online workers should produce chance results. This would be expected when the presentation intensity was appropriate for the source (i.e., low for low-intensity sources, and high for high-intensity sources). The responses might then be biased upwards or downwards as the presentation intensity increased or decreased from this level.

### 3.1. Method

#### 3.1.1. Participants

210 online workers participated in Experiment 1 (113 female; mean age = 33.5 years; SD = 14.7 years) – five for each of the 42 participants.

#### 3.1.1. Materials and procedure

Each text description typed by listeners in Experiment 1 was paired with the original description of the sound (Table S1) and presented to a second set of Mechanical Turk workers. The workers were told that the two descriptions were both provided by listeners, and were asked to choose which of the two sounds being described was the “louder” source. The order in which the two options were presented (source label and listener description) was randomized. Workers were instructed that if the two descriptions were very similar, they should guess. Workers were not given any other information about the sounds or the experiment details. Five online workers independently performed this judgment for each response from a participant in the original experiment. Each data point in the results graph thus reflects 2250 worker scorings (of 450 in-lab typed descriptions). As a consequence, the results graph (proportion judged louder vs. presentation intensity x source group) was stable across splits of the Mechanical Turk responses (split-half Pearson correlation was 0.99).

### 3.2. Results and discussion

The causal inference hypothesis predicts that a source presented at an atypically high intensity will be misidentified as a “louder” source – because the original source could not possibly have created such a high-intensity sound. Thus, the fraction of sources misidentified as “louder” should increase with presentation intensity, and should be higher for low-intensity sources at all presentation intensities (because for any given presentation intensity, more low-intensity sources will be louder than normal than high-intensity sources). The results (Fig. 2F) show that both these effects are observed: the listeners in Experiment 1 were more likely to describe a high-intensity source when the presentation intensity was high (F(6,246)=24.0, p<0.001, 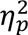 =0.369), and such errors were more common for low-intensity sources (F(1,41)=57.7, p<0.001, 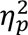 =0.584). In addition, chance performance occurred at low presentation intensities (40-50 dB SPL) for low-intensity sources, and high intensities (90 dB SPL) for high-intensity sources. These results are consistent with the idea that listeners infer the intensity of the source, and give descriptions that are consistent with this inferred intensity.

To assess whether the high-intensity source labels that were mistakenly chosen by participants tended to identify sounds that were otherwise acoustically similar to the presented low-intensity source, we analyzed their acoustics. For each low-intensity source mistakenly identified by a participant, we measured both the mean power in each of a set of gammatone filters (commonly termed the “excitation pattern”), and the mean power in each of a set of spectrotemporal modulation filters (see Appendix E for details of the filter banks). We then made the same power measurements in the high-intensity source recording corresponding to the erroneously selected label, as well as a distinct high-intensity source recording with the closest loudness rating (from Table S1). We then compared the correlation of the power measurements in the low-intensity source with those for each of the two high-intensity sources (after subtracting out the mean of the power measurements across the entire sound set). This analysis revealed that participants tended to choose high-intensity source labels corresponding to sounds whose modulation statistics were more similar to those of the presented low-intensity source than would be expected by chance. Specifically, the correlation with the chosen label (median = 0.33) was significantly higher than that with the unrelated label (median = 0.05) by a sign test (p=0.009). There was no such relationship for the excitation pattern (p=0.33), consistent with the idea that higher-order statistical properties are more important for sound identification than the spectrum (McDermott & Simoncelli, 2011).

## 4. Using Reverberation to Convey Source Distance

Another stimulus variable that should affect inferred source intensity is reverberation. In natural scenes, source-listener distance affects reverberation in a characteristic way. The direct arrival (i.e., the first and highest-intensity peak of the impulse response) decreases in intensity with source-listener distance according to the well-known inverse square law (Fig. 3A). However, the average intensity of reflected sound (i.e., the reverberation) often does not change appreciably with distance because as distance increases, some reflection paths decrease in length and others increase in length. The Direct-to-Reverberant Ratio (DRR), which compares the intensity of the direct sound to that from all the reflections, therefore generally decreases with source distance both indoors (Bronkhorst & Houtgast, 1999; Mershon & Bowers, 1979; Zahorik, Brungart, & Bronkhorst, 2005) and outdoors (Naguib & Wiley, 2001) (Fig. 3A). Once convolved with a source signal, the DRR is no longer explicitly available in the sound signal. However, humans can recognize source distance from reverberant recordings, and are thought to estimate the DRR to do so (Bronkhorst & Houtgast, 1999; Mershon & Bowers, 1979; Zahorik et al., 2005).

**Figure 3.**
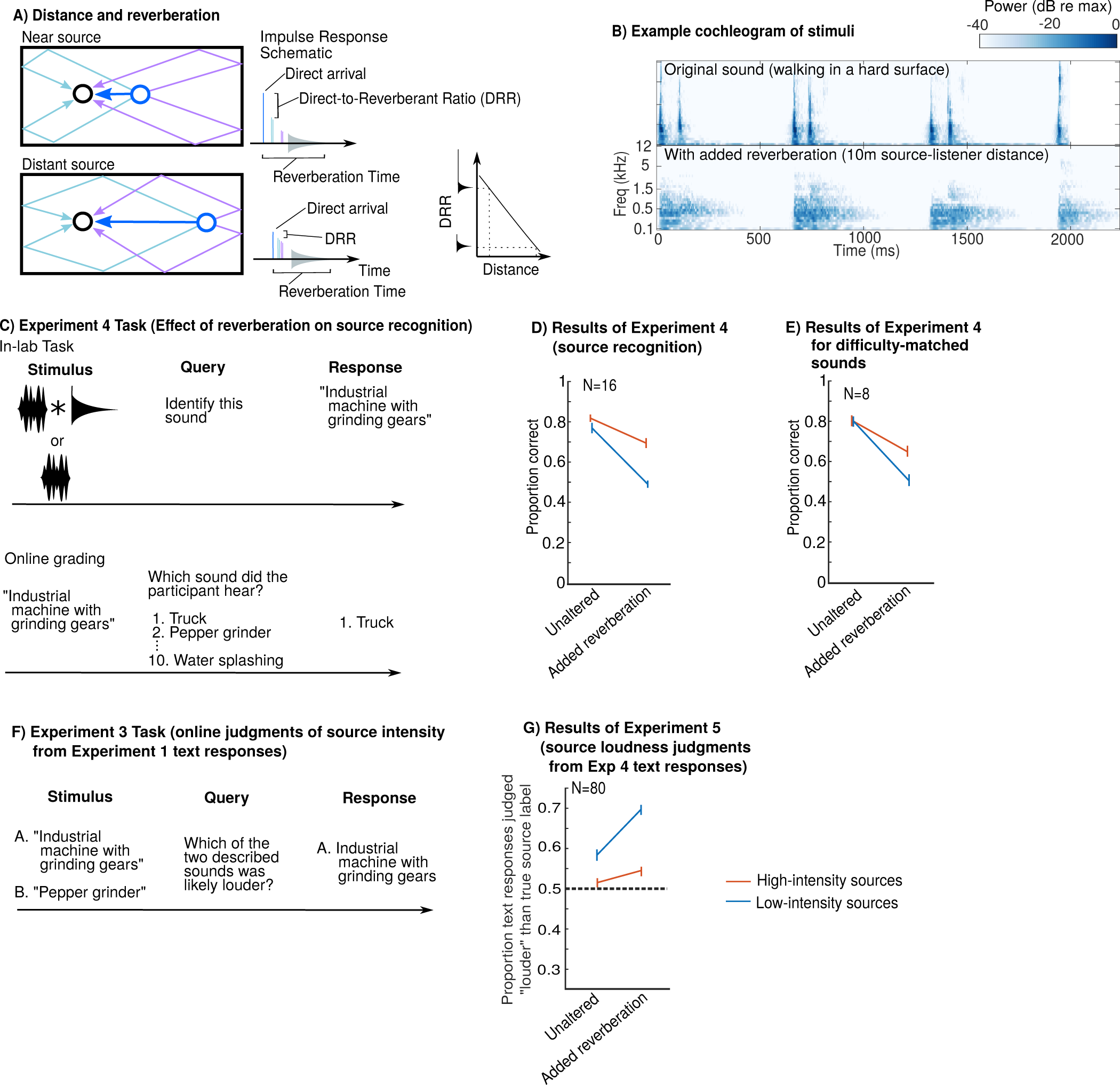
(A) Illustration of the effect of distance on reverberation. When a source is near the listener (top) the direct path (blue) is short (left) and creates a high amplitude peak in the Impulse Response (right). When a source is distant from the listener (bottom), the direct path is longer, producing a correspondingly lower amplitude peak in the Impulse Response. In contrast, the total contribution of the reflections, which arrive after the initial peak, is similar for near and distant sources. We show 2^nd^-order reflections (i.e., the paths that reflect off of 2 surfaces before arrival), which hit both the near and far wall. For simplicity neither 1^st^-order nor higher-order reflections are shown in the room schematic. The acoustic contribution of higher-order reflections is shown in grey at right. Two of the 2^nd^-order reflections (near-wall; green) have longer paths and lower amplitude peaks, but two (far-wall; purple) have shorter paths and are correspondingly louder. Thus, the total contribution from 2^nd^-order reflections is not substantially changed by distance. The same logic applies to all higher-order reflections. Because the power of direct-path sound decreases with distance, the Direct-to-Reverberant ratio (top right) decreases with source distance. Thus, the presence of reverberation with small DRR implies greater distance and thus a more powerful source for a given sound intensity at the ear. Note that impulse responses also vary in the decay time, commonly quantified as the RT60. In simple indoor conditions, the DRR and RT60 can vary independently. The RT60 is typically fairly constant within a room, and varies across rooms depending on their size and on the material of their walls. By contrast, the DRR varies within a room depending on the distance of the source to the listener. (B) Cochleagrams of an example environmental sound from Experiment 5 (the sound of walking) without (top) and with (bottom) added reverberation. The reverberation was synthesized to be typical of a 10m separation in a large room. (C) Schematic of Experiment 4 (effect of reverberation on source recognition). (D) Results of Experiment 4: Recognition declined more for low-intensity than high-intensity sources when presented in reverberation, consistent with causal inference of source intensity. (E) Results of Experiment 4 with difficulty-matched subsets of sources. (F) Schematic of Experiment 5 task (identical to Experiment 3). (G) Results of Experiment 5: The fraction of typed responses from Experiment 4 which were judged to correspond to louder sources than the true source label, as a function of the reverberation and source-intensity.

## 5. Experiment 4: Reverberation Impairs Recognition of Low-Intensity Sources

In Experiment 4, we asked participants to identify sound sources both with and without the addition of synthetic reverberation that implied a distant source. Under the causal inference hypothesis, greater implied distances via reverberation should be used to infer greater source intensities, and should produce similar errors as Experiments 1-2 even when intensity at the ear is held constant. Specifically, low-intensity sources should be misidentified at greater rates than high-intensity sources when rendered at a distance. By contrast, under an invariance hypothesis, recognition should be dependent on the extent to which reverberation could be separated from the sound source. This need not entail complete invariance to reverberation, but whatever invariance might be achieved should be similar for low- and high-intensity sources.

### 5.1. Method

#### 5.1.1 Participants

16 in-lab listeners (7 female, 9 male; mean age = 40.1 years; SD = 13.5 years) took part. Pilot experiment data (not included in the results presented here) suggested an interaction effect size of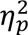=0.481. A power analysis indicated 11 participants were needed to detect an effect of this size 80% of the time using a significance threshold of .05.

#### 5.1.2 Stimuli and Procedure

All experiments manipulating reverberation (Experiments 4-11) used a subset of 192 sounds (Table S1). This subset contained neither speech nor music but was otherwise a representative and randomized subsampling of the original 300. The intensity- and reverberation-experiments were originally begun as separate studies and thus were not designed to match exactly. In all reverberation-experiments (Experiments 4-11) “high/low-intensity sounds” refers to the upper/lower halves of the 192-sound subset (i.e., 96 sounds) rather than the upper/lower quartiles, as in the intensity experiments (1-3 and 12).

Each listener heard each sound once, presented either with or without reverberation. The reverberation conditions were balanced across listeners, such that while each listener only heard each sound once, each source was equally likely to be presented reverberant or anechoic.

We intended for our reverberation manipulations to exceed the magnitude of any incidental reverberation in the recordings, and so applied fairly pronounced synthetic reverberation. The reverberation (used in Experiments 4-11) was synthesized as described in (Traer & McDermott, 2016) with a Direct-to-Reverberant Ratio (DRR) of 20dB, consistent with a source-receiver separation of about 10m. We gave the reverberation a broadband decay time (RT60) of 1s, consistent with a large interior space such as a subway station, such that the overall amount of reverberant energy in the resulting sound signal was high. Fig. 3B shows cochleagrams of an example stimulus with and without added reverberation.

The in-lab task and online scoring were identical to Experiment 1 (Fig 3C) except that all the sounds were presented at 70dB SPL and each listener’s response was graded by 5 different workers instead of 2.

### 5.2 Results and discussion

As shown in Fig. 3D, recognition was worse overall in reverberation, as expected given the substantial distortion imposed by the reverberation (main effect of reverberation: F(1,15)=252, P<0.001, 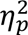=0.944). However, the effect of the reverberation was less pronounced for high-intensity sources, producing a significant interaction between reverberation and source intensity (F(1,15)=62.0, p<0.001, 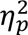 =0.805). To ensure the interaction was not driven by differences in difficulty between the source classes, we equated overall recognition rates for the condition without added reverberation, by eliminating the most-recognized and least- recognized sources when presented without reverberation in each group. The matched sets were obtained with data from half of the participants, and the data from the other half is plotted in Fig. 3E. As is evident in Fig. 3E, the interaction between the effect of presentation intensity and the source intensity persisted after this manipulation (F(1,7)=16.7, p=0.005, 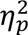=0.705).

## 6. Experiment 5: Recognition Errors in Reverberation Support Causal Inference

As an additional test of the causal inference hypothesis, we assessed whether listeners exhibited the pattern of errors predicted by causal inference, tending to misidentify a low-intensity source in reverberation as a high-intensity source. The text responses of Experiment 4 were graded as in Experiment 3, with online graders judging whether the written responses described a sound that was louder or quieter than the actual sound listeners heard. Because the sounds were presented at 70 dB SPL, the causal inference hypothesis predicts that reverberant low-intensity sources will be misidentified as “louder” sources (replicating the effect of Experiment 3). The 10m source distance implied by the added reverberation might increase this effect (because increased distance should imply a higher-intensity source for a fixed presentation intensity at the ear). By contrast, high-intensity sources are less likely to be affected by reverberation in this way, as they are not inconsistent with the intensity implied by distance.

### 6.1 Participants

80 online workers participated in Experiment 5 (44 female; mean age = 37.2 years; SD = 16.3 years). 5 workers graded each text response collected in Experiment 4 (the same number used in Experiment 3).

### 6.2 Method

Experiment 5 was identical to Experiment 3 (Fig 3F), except that it was performed on the text responses from Experiment 5 (reverberation manipulation), rather than those of Experiment 1 (intensity manipulation).

### 6.3 Results and discussion

As shown in Fig. 3G, there was an overall tendency for written descriptions of low-intensity sources to suggest higher-intensity sources than the original source labels (t-test against chance: t(15) = 49.5, p<0.001), with no such effect for the high-intensity sources. This result without the added reverberation replicates the effect of Experiment 3 for the 70dB condition. Moreover, as predicted by causal inference, there was an interaction with reverberation (F(1,15)=28.4, P<0.001, 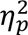 =0.654): for low-intensity sources there was larger difference between the conditions with and without reverberation (t(15) = 8.95, p<0.001, Cohen’s D = 2.71)), than for high-intensity sources (t(15) = 3.56, p=0.003, Cohen’s D = 0.802). This result supports a causal inference interpretation of Experiment 4: it appears that reverberation increases perceived distance which in turn increases the inferred source intensity, causing systematic errors for the low-intensity sources.

## 7. Experiment 6: Effect of Reverberation on Perceived Distance in Natural Recordings

In Experiments 4-5 we found that reverberation impaired recognition of low-intensity sources more than high-intensity sources, plausibly because it implies distance, and thus implies a higher-intensity source for a given intensity at the ear. To further assess this explanation, we evaluated the distance attributed to the sound sources in our stimuli.

Measuring perceived distance seemed particularly important given our use of real-world recordings. The use of such recordings enabled a large and diverse stimulus set, but comes at the cost of occasional background noise and unavoidable reverberation. As a consequence, the recordings used in Experiments 1-5 all had some reverberation from the space in which they were recorded. At present, there is no available method to quantify such reverberation from a recording. However, we can instead assess the perceptual effect of potential reverberation by having participants estimate the distance of the sound sources.

Because this experiment was quite short in duration, it was conducted online. Our lab has previously found that listening experiments run online generally replicate data collected in the lab, qualitatively and quantitatively (McPherson et al., 2020; McPherson & McDermott, 2020; McWalter & McDermott, 2019; Woods & McDermott, 2018), provided steps are taken to ensure participants comply with instructions (McPherson & McDermott, 2020; Woods, Siegel, Traer, & McDermott, 2017).

### 7.1. Method

#### 7.1.1 Participants

80 online listeners participated in Experiment 6 (36 female, 42 male, 2 did not report; mean age = 43.2 years; SD = 10.01 years) via Amazon’s Mechanical Turk. We had no pilot data with which to run an a priori pilot analysis, but data collection was fast and inexpensive, so we ran a relatively large number of online participants to err on the side of being over-powered. All online listeners in this and other experiments in this paper self-reported normal hearing. All online listening tasks included a test at the start of the experiment to help ensure that listeners were wearing headphones (Woods et al., 2017). The participants analyzed and reported for each online experiment all passed this test.

#### 7.1.2 Stimuli and Procedure

The stimuli and procedure for Experiment 6 were identical to that of Experiment 4 (same sounds, reverberation, and balancing of classes), except instead of describing sounds, listeners were asked to guess the distance of the sound source from microphone (Fig. 4A). There were seven logarithmically-spaced response options: 10cm (4 inches); 30cm (1 ft); 1m (3 ft); 3m (10 ft); 10m (30ft); 30m (100ft); 100m (300ft). Listeners were not given any other information (e.g., the source identity) and they were told in advance that all sounds were artificially constrained to have the same intensity level, such that intensity was not a reliable cue to distance.

**Figure 4.**
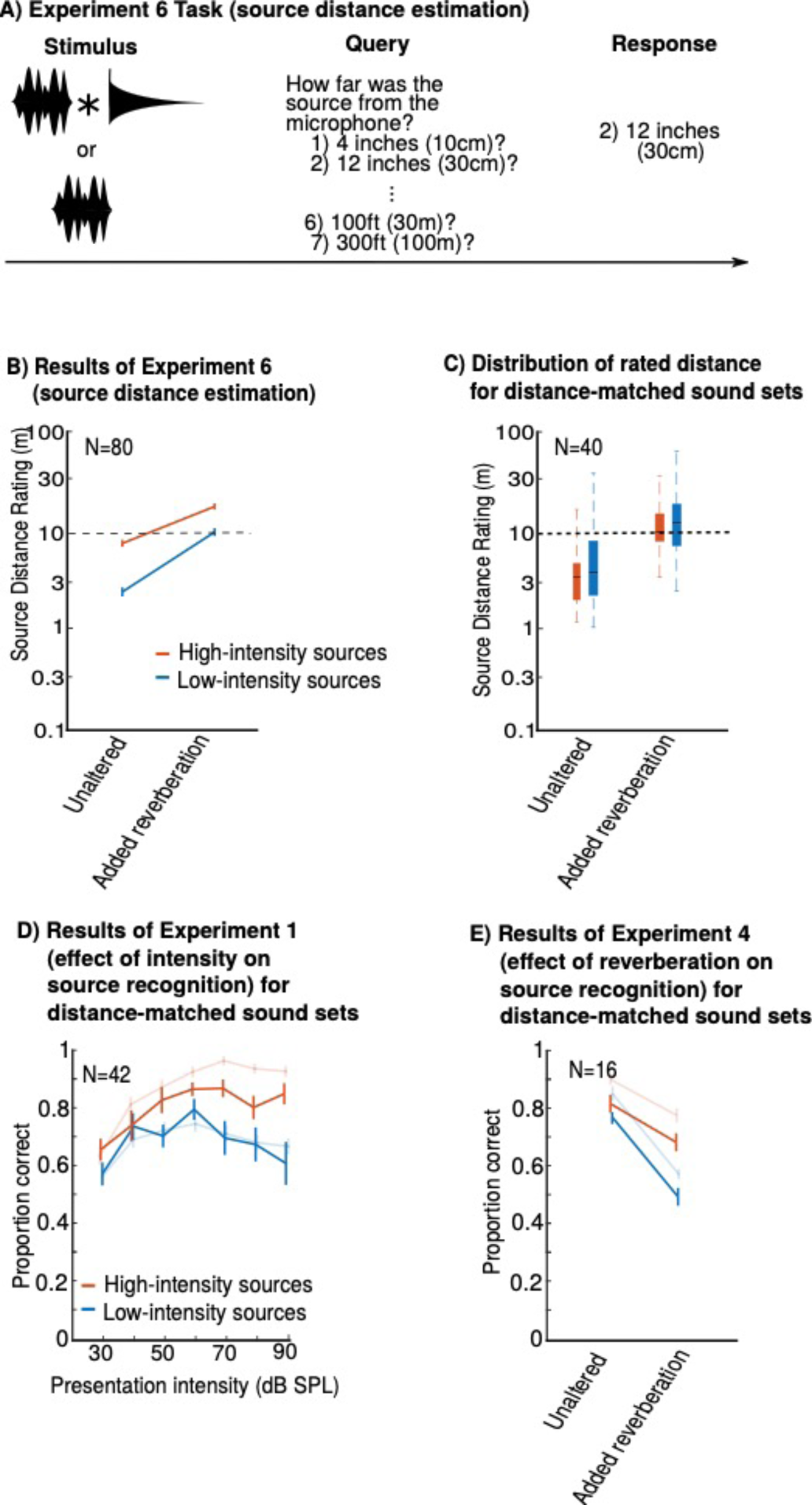
Natural reverberation affects the perceived distance of the stimuli, but does not drive the source recognition effects from Figures 2 and 3. (A) Schematic of Experiment 6: online listeners heard sounds with and without added reverberation and estimated the distance between the source and microphone. (B) Results of Experiment 6: Judged distance for sounds with and without reverberation, plotted separately for high- and low-intensity sources. The dashed line shows the distance that synthetic reverberation was designed to emulate (10m). For all sources, adding synthetic reverberation increased perceived distance. However, high-intensity sources were judged as more distant than low-intensity sources, likely due to reverberation in the original recordings. The dashed line shows the distance that synthetic reverberation was designed to emulate (10m). (C) The distribution of distance ratings from Experiment 6 for “distance-matched” subsets of the stimuli. The subsets were chosen by eliminating the most distant high-intensity and the least distant low-intensity sources. Data from half of the Experiment 6 participants were used to choose the sounds, and data from the other half are plotted here. (D-E) The results of Experiment 1 (D) and Experiment 4 (E) with analysis restricted to the distance-matched subsets. The results are similar to those for the full sets of sounds (shown transparent), indicating that differences in the perceived source distance between the two sets of sounds do not account for the recognition differences.

Due to the constraints of running the experiment online, we could not control the absolute presentation level of the stimuli, but all stimuli had the same rms level, and participants were instructed to adjust their volume setting using a calibration sound such that the experimental stimuli were comfortably audible.

### 7.2 Results and discussion

As shown in Fig. 4B, the added reverberation increased the perceived distance of the sound source in all cases, as intended, but there were also pronounced differences between sound categories. Specifically, high-intensity sources were judged to be further away than low-intensity sources. This likely reflects practical constraints on sound recording, whereby high-intensity sound sources (e.g., a truck backing up, freight train, etc.) must be recorded at a distance, with concomitant reverberation cues in the recorded sound. By contrast, low-intensity sources are often recorded in quiet environments with a close microphone. Two other factors could also contribute. First, if listeners use knowledge of typical source intensities to calibrate distance judgments, high-intensity sources would be expected to seem further away, all other things being equal (as they were here, with all sounds presented at the same intensity). Second, listeners could plausibly have learned source-distance associations (e.g., because high-intensity sources might be more often encountered at far distances) and might be influenced by them when estimating distance, thus judging the high-intensity sources to be further away irrespective of reverberation.

The distance cues in the original recordings remained present when synthetic reverberation was added: the synthetic reverberation was designed to simulate a 10m source-microphone separation in a large room, and although judged distances were in the neighborhood of this value (between 5-20m), the high-intensity sounds were judged as substantially more distant than the low-intensity sounds even with added synthetic reverberation.

These results are consistent with the possibility that the reverberation present in typical real-world recordings interferes with the ability to manipulate perceived distance with reverberation.

Given this, it seemed important both to attempt to control for the differences in distance in our recordings, and to make more controlled recordings in which distance cues would be minimal (and fixed across the sounds to be compared).

## 8. Distance-matched sound sets

Given the variation in perceived distance across the stimuli in the absence of added reverberation, we sought to use “distance-matched” subsets of high-intensity and low-intensity sources with which we could test the effect of distance on source recognition in a more controlled manner. From the original sets of sounds used in Experiments 4-6, the most distant high-intensity source and the least distant low-intensity source were iteratively excluded until the distributions overlapped (i.e., until the difference in the mean rated distance for the two groups was less than 0.1 on the 1-7 scale used in the rating, corresponding to a distance ratio of 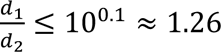 of the two mean distance ratings, *d*_1_ and *d*_2_). This selection procedure was performed using data from half the participants of Experiment 6 (N=40). To verify the success of the procedure, we used the data from the other half of participants to measure the average perceived distance of the resulting groups of sounds (Fig. 4C)). This yielded distance-matched subsets of 26 high- and 26 low-intensity sounds (Table S2).

**Table 2:**
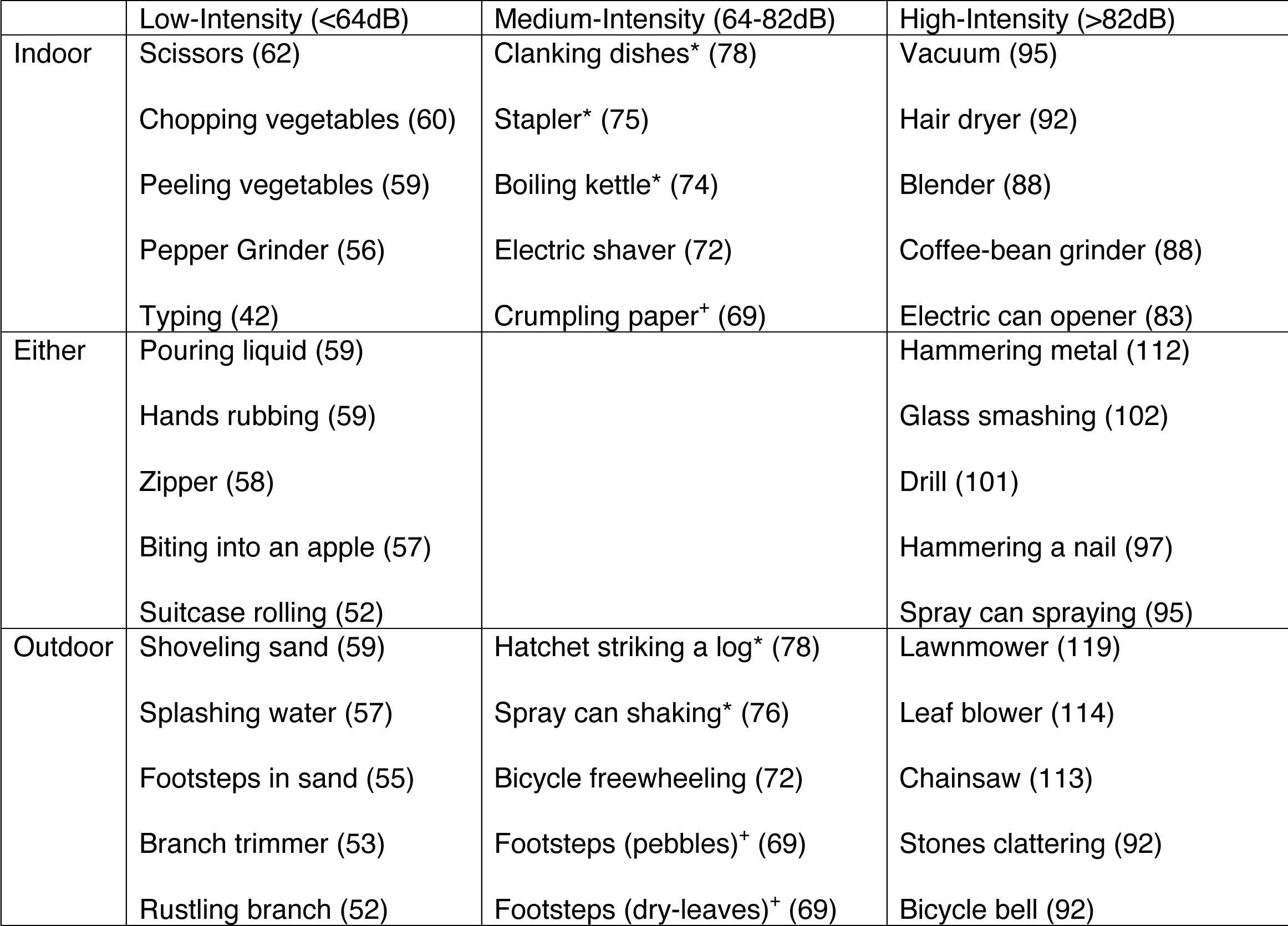
Sounds recorded in a studio with minimal reverberation. The sound level (in dB SPL) as measured 10cm from each source is given in parentheses. In comparisons of high-intensity vs. low-intensity sources (Fig. 5) the 15 sounds from the left- and right-columns are used. In comparisons of indoor vs. outdoor sources (Fig 6) the 15 sounds from the top and bottom rows are used. The Medium-Intensity sources marked with an asterisk or dagger were classed as high- or low-intensity sources, respectively, in the analyses shown in Figure 6 to increase the pool of sources.

The distribution of distance ratings for these subsets of recordings are shown in Fig 4C, with and without synthetic reverberation. Without added reverberation, the rated distance was matched across source types, as intended. With synthetic reverberation, perceived distance increased, as intended, and to a similar extent across source types. Moreover, distance ratings with the added reverberation were close to 10m, demonstrating a reasonable quantitative match between intended and perceived distance. These results indicate that we achieved the desired manipulation of perceived distance.

To assess whether the key results were robust to the incidental distance cues present in the sound recordings, we replotted the results of Experiments 1 and 4 including only the distance-matched subsets (Fig 4D-E). As we found with the full set of sounds, source-recognition was impaired when low-intensity sources were presented at high intensities (again producing a significant interaction: F(2,82)=4.09; p=0.020; 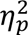=0.091; because the number of sounds was reduced we binned the presentation intensity groups into three bins (less than 50dB; 50— 70dB; greater than 70dB) to ensure all participants encountered at least 3 sounds per condition (average of 7.4)). Source recognition for low-intensity sources was also impaired by reverberation, again producing a significant interaction (F(1,15)=16.0, p=0.001, 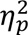 =0.516). These results suggest that the variation in apparent source distances present in the original stimulus set cannot explain the different effects of intensity at the ear and reverberation on the recognition of high- and low-intensity sound sources.

## 9. Studio recordings with controlled reverberation

Although our main results were reproduced with distance-matched subsets of sounds (Fig 4), the distance ratings (Experiment 6) showed large and systematic differences between the perceived distance of different types of sound, probably due to differences in reverberation contaminating the original recordings. Given that the results of Experiments 4 and 5 suggest that reverberation affects source recognition, we sought to replicate our main findings with an additional set of sounds recorded in a soundproof studio to minimize reverberation (see Appendix D). The studio had damped walls and we used a fixed small (10cm) source- microphone distance (Fig 5A). The sound sources were chosen to span different source intensities (15 each of high- and low-intensity sources; see Table 2 and Fig 5B) and to include both indoor and outdoor sound sources, as this distinction was important for Experiment 11. In addition to recording the sound we measured its sound pressure level at the microphone, using a sound level meter. These controlled recordings allowed us to more carefully test the effect of reverberation on perceived source distance (Experiment 7) and source recognition (Experiments 8 and 9).

**Figure 5.**
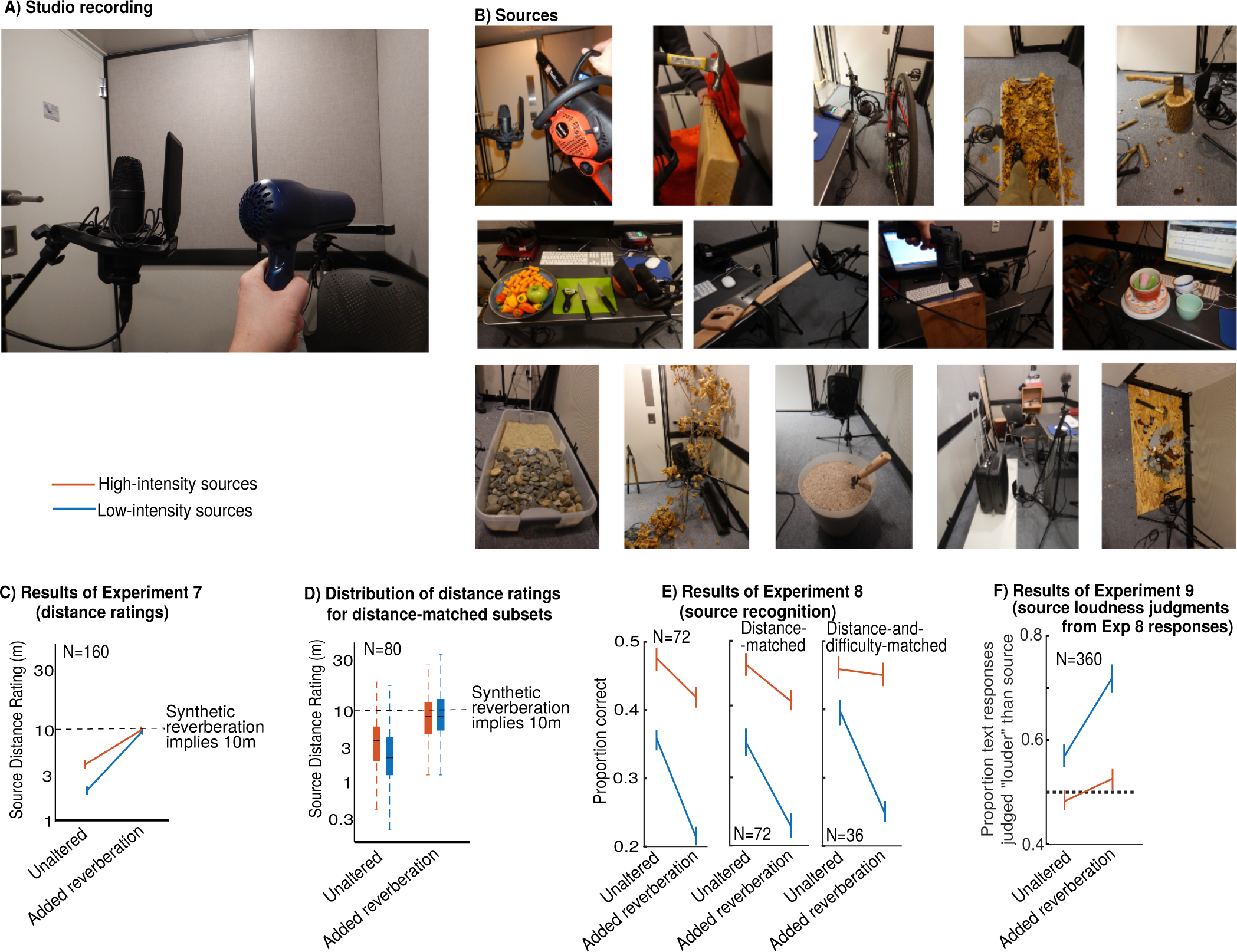
Studio-recorded sources with minimal reverberation. (A) Recording environment: sources were recorded in an acoustically damped sound booth with a 10cm source-microphone spacing. The image shows the setup for recording a hair-dryer. (B) Example recorded sources (from top-left): chainsaw, hammering nails into wood, bicycle freewheeling, walking in dry leaves, chopping wood with a hatchet, chopping and peeling vegetables, sawing wood, an electric drill, clattering of dishes, walking on stones and sand, rustling branches, shoveling sand, wheeling a suitcase, and glass shattering. (C) Results of Experiment 7: Judged distance for sounds with and without reverb broken down by whether the source intensity was low or high. (D) The distribution of distance ratings from Experiment 7 for two “distance-matched” subsets of the natural recordings (data that are plotted are distinct from those used to choose the subsets). (E) Results of Experiment 8 (source recognition of studio-recorded sources) with all data (left), distance-matched sounds (middle), and sounds matched in both distance and difficulty (right). (F) Results of Experiment 9 (Loudness judgments of the written descriptions from Experiment 8. Analysis was restricted to the distance-matched subsets. In both E and F, the interaction effects are similar to those of the natural recordings, suggesting that the interactions in Experiments 1-5 are not driven by contaminant reverberation.

## 10. Experiment 7: Effect of Reverberation on Perceived Distance of Studio-Recorded Sources

To test the effect of added reverberation on perceived source distance for our studio-recorded sounds, we repeated Experiment 6 (distance estimation) but with the new set of recorded sounds (Fig 6C).

**Figure 6.**
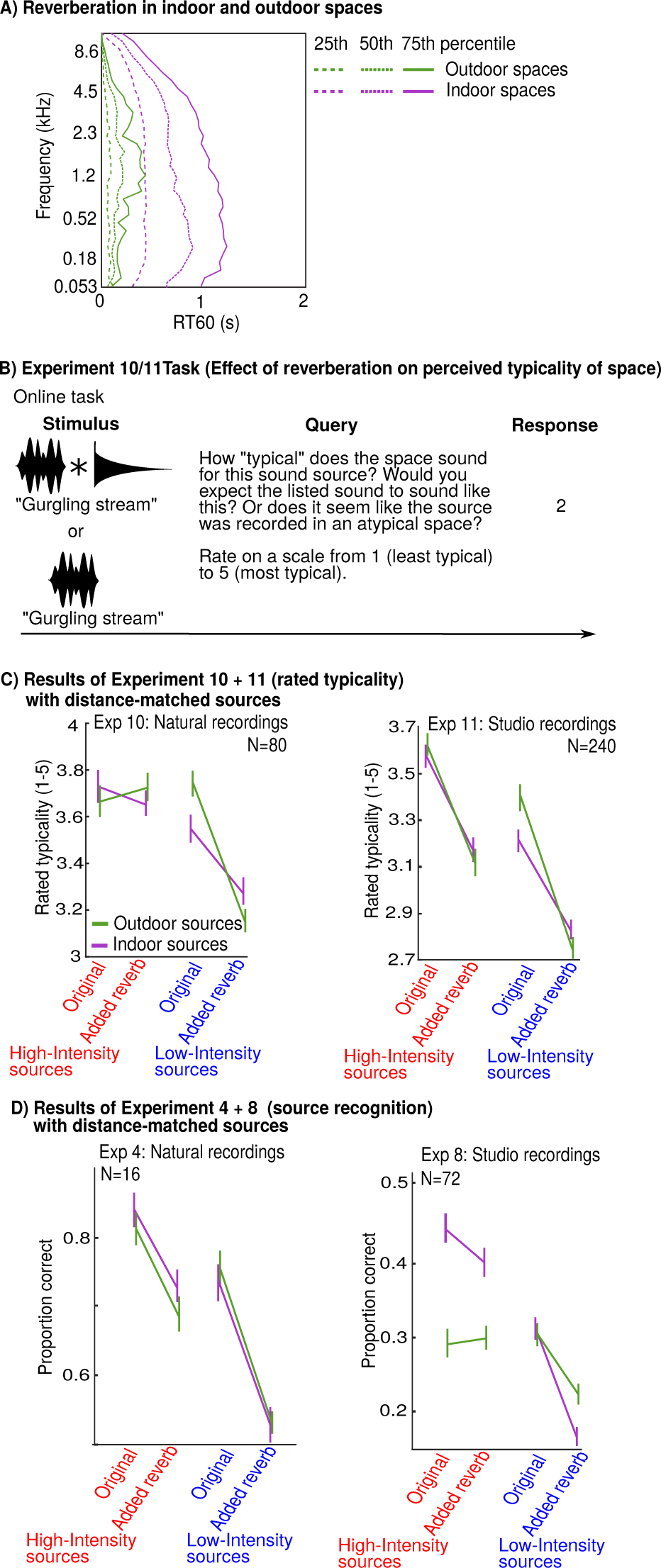
Interactions between reverberation and typical source location support causal inference. (A) Typical frequency-dependent reverberation times (RT60) for indoor and outdoor spaces, measured with a 2m source- microphone separation. Measurements were obtained in a survey of ecological reverberation (data replotted from Traer and McDermott, 2016) and the three lines for each location group show the 25^th^, 50^th^, and 75^th^ percentiles of RT60 at different frequencies. Thus, for equivalent source-listener distances, indoor sounds would be more commonly encountered with reverberation than outdoor. (B) Schematic of the task used in Experiments 10 and 11, in which participants judged the typicality of a sound’s audible environment with and without application of synthetic reverberation, for natural recordings (Experiment 10) or studio recordings (Experiment 11). (C) The results of Experiments 10 and 11: low-intensity outdoor sources suffered a greater decrement in rated typicality of their environment when reverberation was applied than low-intensity indoor sources. The difference between indoor and outdoor sounds was not observed for high-intensity sounds, plausibly because loud sources can be heard from great distances, and over a large enough distance outdoor environments can exhibit significant reverberation. (D) Results of Experiments 4 and 8 (source recognition) with the results plotted separately for indoor and outdoor sources. There is no evidence that reverberation impairs recognition more for low-intensity outdoor sources than low-intensity indoor sources, despite the difference in the appropriateness of the reverberation observed in Experiments 10 and 11. This suggests that misidentification of sources is not being driven by the atypicality of the reverberation, but rather is caused by the physical implausibility of a reverberant low-intensity source.

### 10.1. Method

#### 10.1.1 Participants

160 online listeners participated in Experiment 7 (92 female, 7 did not report; mean age = 29.4 years; SD = 9.00 years). We ran twice as many participants as in Experiment 6 (192 sounds), because there were fewer sounds in this experiment (45 studio recordings, along with 72 natural recordings as controls to ensure performance was comparable to that of in-lab participants).

#### 10.1.2 Stimuli and Procedure

The experiment was identical to Experiment 5 except that the natural recordings (Table S1) were replaced with a set of studio recordings (Table 2), with a sampling rate of 44.1 kHz and a bit depth of 24.

### 10.2 Results and discussion

As expected, the studio recordings (before synthetic reverberation was added) were rated overall as less distant than the natural sound sources of Experiments 1-6 (2.84m vs. 4.86m, on average). And as intended, the synthetic reverberation increased distance judgments to approximately 10m. In addition, the difference in perceived distance between high- vs. low-intensity sources without added reverberation was much smaller for the studio recordings than for the natural recordings used in Experiment 6 (1.80m vs. 6.27m). This difference suggests that the large differences between the source types in Experiment 6 were in part driven by reverberation in the natural recordings. However, high-intensity sources were nonetheless rated as more distant than low-intensity sources (t(159)=5.19, p<0.001, paired t-test). Moreover, the distance estimates without added reverberation consistently exceeded the actual source-microphone distance of 10 cm (t(159)=66.4; p<0.001, t-test vs. 10cm). These differences suggest a role for the additional factors noted earlier. Given that the sounds were all presented at the same intensity, distance could be calibrated by knowledge of typical source intensities, causing high-intensity sources to seem further away. Alternatively, the difference between high- and low-intensity sources could reflect learned source-specific priors on distance.

Overall, these results indicate that the distance manipulation largely works as expected when applied to recordings with minimal reverberation. But given that there were still small differences in rated distance between the different sound classes before reverberation was added, we ran the distance matching procedure again, yielding subsets of 13 high- and low-intensity sounds (Fig 5D; Table S4).

## 11. Experiments 8 and 9: Effect of Reverberation on Recognition of Studio-Recorded Sources

To confirm the source-recognition results of Experiments 4-5, we replicated both experiments with the distance-matched subsets of studio-recordings. Because the set of studio recordings was much smaller than the set of natural recordings, Experiment 8 was run online to obtain a large number of participants and thus sufficient power.

### 11.1. Method

#### 11.1.1. Participants

72 online listeners participated in Experiment 8 (37 female, 34 male, 1 did not report; mean age=31.25; SD = 14.28). Pilot experiment data (not included in the results here) suggested an interaction with effect size of 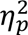=0.109. A power analysis indicated that 67 participants were needed to detect an effect of this size 80% of the time using a significance threshold of .05.

360 online workers participated in Experiment 9 (174 female, 8 did not report; mean age = 33.2 years; SD = 16.8 years). This number resulted in 5 workers grading each text response of Experiment 8, the same number as in Experiments 3 and 5.

#### 11.1.2. Materials and procedure

Experiments 8 and 9 were identical to Experiments 4 and 5, except that studio-recorded sources were used instead of natural recordings. See Table S3 for a list of distance-matched studio-recordings. The grading procedure was also identical to that used in Experiments 4 and 5.

### 11.2 Results and discussion

As shown in Fig. 5E, recognition was again worse overall in reverberation, as expected given the substantial distortion imposed by the reverberation (main effect of reverberation: F(1,71)=48.44, P<0.001, 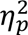 =0.406). However, the effect of the reverberation was less pronounced for high-intensity sources, producing a significant interaction between reverberation and source intensity (F(1,71)=8.85, p=0.004, 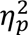 =0.111), replicating the effect observed in Experiment 4.

The lower overall performance for the studio recordings compared to the natural recordings appears to reflect idiosyncrasies of the set of sources (chosen based on practical constraints of being able to record them in a small studio). It could in principle reflect differences between in-lab and online performance, but the online participants also performed a small number of “sanity-check” trials with a subset of the natural recordings. For these trials, their overall mean performance was 71%, which was comparable to that of the in-lab participants (69%). This suggests the studio recordings are intrinsically more difficult to recognize and describe than the natural recordings. We note that the low recognition rates are still well above chance (10%).

It was also the case that the high-intensity studio sources were overall more recognizable than the low-intensity studio sources. To control for this difference we selected subsets of 8 high- and 8 low-intensity sources that were matched for both recognizability and distance without reverberation. The difficulty-matched sources were selected using the data from half the participants and the data from the other half is plotted in 4E. Because of the small number of stimuli, it was not possible to perfectly equate recognizability in this way, but the difference between conditions was substantially reduced compared to the main analysis. These difficulty-matched subsets still showed a significant interaction between source intensity and reverberation (F(1,35)=5.43, p=0.026, 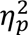=0.134).

Experiment 9 replicated Experiment 5 but with the descriptions of the distance-matched subset of studio recordings. As shown in Fig. 5F, the written descriptions of low-intensity sources presented in reverberation again suggested higher-intensity sources than the original source labels (t-test: t(35)=9.00, p<0.001, Cohen’s D = 1.45). This effect was again absent for high-intensity sources (t-test: t(35)=0.966, p=0.341, Cohen’s D = 0.178), producing an interaction between source intensity and reverberation (F(1,35)=9.19, p=0.005, 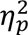=0.208), replicating the effects observed in Experiment 5 and again providing further support for the causal inference interpretation.

## 12. Causal Inference vs. Acoustic Familiarity

The results of Experiments 1-9 are inconsistent with the invariance hypothesis, and are consistent with the idea that listeners use inferred source intensity as a recognition cue. We refer to this possibility as causal inference. However, the key result – that low-intensity sources are misidentified when presented at unusually high-intensities or distances – is consistent with at least one other hypothesis: that recognition is constrained by whether a listener has previously heard a source in the presentation conditions (as might be expected if listeners learn a set of templates of their previous sensory experience, and recognize sounds via matches with these templates). The causal inference hypothesis could explain the results given that neither high-intensity nor reverberant sounds could be caused by a low-intensity source. The acoustic familiarity hypothesis could also explain the results because low-intensity sources would never be encountered as such in natural scenes.

To distinguish these two hypotheses, we reanalyzed the data from the source-recognition experiments in reverberation (Experiments 4 and 8) and compared the effect of reverberation on recognition of sources typically encountered outdoors against those typically encountered indoors. Although extremely distant outdoor sources can be very reverberant (e.g., thunder, fireworks, or distant gunshots), it is plausible that for sounds of moderate intensity, which are only audible at moderate distances, outdoor scenes would yield shorter reverberation decay times than indoor scenes, as shown for empirical measurements in Fig 6A (data from (Traer & McDermott, 2016)). The shorter decay times reduce the overall amount of reverberant sound energy in the signal reaching the ears, all other things being equal. Under the acoustic familiarity hypothesis, low-intensity outdoor sounds should be more often misidentified than low-intensity indoor sounds when our synthetic reverberation is applied, whereas under the causal inference hypothesis, there should be no difference.

## 13. Experiments 10 and 11: Reverberation is Unnatural for Outdoor Sources

Before re-analyzing the recognition results, we conducted an experiment to test whether added reverberation would be heard as “less typical” for outdoor compared to indoor sound sources, and thus test the key assumption motivating the re-analysis (Fig 6A). As with Experiments 6 and 7, this experiment was short in duration and was thus conducted online. Participants were presented with an audio recording and its label, and were asked to rate, on a scale from 1 to 5, how typical the environment seemed for the named sound source. Participants heard each sound only once, with half the sounds presented unaltered and the other half with added synthetic reverberation. The reverberation had a direct-to-reverberant ratio (DRR) appropriate for a 10m distance, and a long decay time (RT60) appropriate for a large room, as in Experiment 4. Under the assumption that outdoor spaces do not exhibit such long RT60s, as has been demonstrated for nearby sources (Traer & McDermott, 2016), this reverberation is inappropriate for outdoor sounds. Experiment 10 used the natural recordings of Experiments 1- 6, while Experiment 11 used the studio recordings of Experiment 7-9. We separately analyzed the results for high- and low-intensity sources because the assumption that the synthetic reverberation is incongruous with outdoor scenes is less obviously justified for distant sources, which must necessarily be high-intensity.

### 13.1 Methods

#### 13.1.1 Participants

80 online listeners participated in Experiment 10 (43 female, 36 male, 1 did not report; mean age = 28.2 years; SD = 15.43 years). 240 online listeners participated in Experiment 8 (111 female, 122 male, 7 did not report; mean age = 32.2 years; SD = 12.09 years). As with Experiments 5 and 6, we had no pilot data for an a priori power analysis but the experiment was fast and inexpensive, and so a large number of participants were run to err on the side of being over- rather than under-powered. More participants were run in Experiment 11 because the experiment contained fewer sounds than Experiment 10.

#### 13.1.2 Stimuli and Procedure

The sound set in Experiment 10 was the distance-matched set of natural recordings, with each source categorized as indoor/outdoor as well as high-/low-intensity (Table S4; Appendix A). The sound set in Experiment 11 were the distance-matched indoor and outdoor studio- recordings (Table S5).

In both experiments, each listener heard each sound once, presented either with or without reverberation. The reverberation conditions were balanced across listeners, such that while each listener only heard each sound once, each source was equally likely to be presented reverberant or anechoic.

Listeners were shown the name of each source and were given the following instructions:

*How “typical” does the space sound for this sound source? Would you expect the listed sound to sound like this? Or does it seem like the source was recorded in an atypical place? Rate on a scale from 1 (least typical) to 5 (most typical)*.

### 13.2 Results and discussion

As shown in Fig 6C, the results differed for high-and low-intensity sources. Low-intensity sources with added reverberation showed the expected interaction. The sounds were overall heard as less natural in reverberation, but the rated typicality decreased more for outdoor than indoor sounds, as predicted, producing a significant interaction between sound class and reverberation for both experiments (Experiment 10, natural-recordings: F(1,79)=11.9; P=0.001; 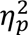=0.0131; Experiment 11, studio-recordings: F(1,239)=7.06, P=0.008, 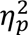=0.029). These effects were not observed for high-intensity sounds, which were rated as about equally typical with or without reverberation for both indoor and outdoor sources (no significant effect of reverberation in Experiment 10, natural-recordings: F(1,79)=0.087; P=0.366; 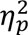=0.011; No significant interaction in Experiment 11, studio-recordings: F(1,239)=0.580, P=0.580, 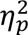=0.002). One explanation is that outdoor environments can exhibit substantial reverberant energy provided a source is sufficiently high-intensity and far away (e.g., a jack-hammer heard from a neighboring street through a window) (Padgham, 2004; Wiener, Malme, & Gogos, 1965). Our reverberation had a decay time that was atypical for nearby sources (e.g., several meters away) in outdoor spaces (Traer & McDermott, 2016), but the reverberation of distant sources in outdoor spaces (Knudsen, 1946) is less well characterized and it is possible that our reverberation is consistent with sufficiently distant sources in outdoor spaces.

Overall, these results provide support for the idea that listeners have some degree of implicit knowledge of the reverberation that is typical for a sound source, such that low-intensity sources can be divided into subsets of outdoor and indoor recordings that might be used to distinguish familiarity-based recognition from causal inference. For low-intensity sounds, which are only audible when close, reverberation is less commonly associated with outdoor than indoor sounds, even though it is no less physically possible.

## 14. Sound Recognition Results (Experiments 4 and 8) Support Causal Inference

When reanalyzed separately for typically indoor and outdoor sounds, the results from Experiments 4 and 8 showed no evidence that low-intensity outdoor sounds were misidentified more in reverberation than indoor sounds (Fig 6C; Experiment 4, natural-recordings (Table S4): F(1,15)=0.127, p=0.727, 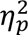=0.008; Experiment 8, studio-recordings (Table S5) showed a significant interaction in the opposite direction: F(1,71)=4.22, p=0.044, 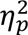=0.056), even though the reverberation we applied was heard as more atypical for the outdoor than indoor sounds in Experiments 10-11. There was also no interaction for the high-intensity sources (Experiment 4, natural-recordings: F(1,15)=0.115, p=0.739, 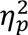 =0.008; Experiment 8, studio-recordings: F(1,71)=1.97, p=0.164, 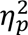=0.027), though none was expected given the results of Experiments 10 and 11.

Overall, these results are consistent with the idea that humans are using implicit causal inference to interpret and identify sources, with the distance cue from reverberation being used to infer source intensity, which then influences recognition judgments.

## 15. Evidence for Causal Inference is Robust to Variations in Choice of Sound Sources

Here, and in Section 16, we present additional analyses and a control experiment to address various alternative explanations of our key results.

We first examined whether the different results for the two groups of sources could be explained by differences in standard acoustic properties (see Appendix E for more details). For each sound, we computed a “cochleagram”, which is similar to a spectrogram but is computed using a filter bank designed to mimic cochlear frequency tuning. We then compared the average spectral power distribution from this filter bank (Fig. 7A) (also known as the excitation pattern, obtained by averaging the cochleagram across time) as well as the power in a set of modulation filters that measure the strength of fluctuations in the cochleagram across time and frequency (Fig. 7B) (Chi, Ru, & Shamma, 2005; Singh & Theunissen, 2003).

**Figure 7.**
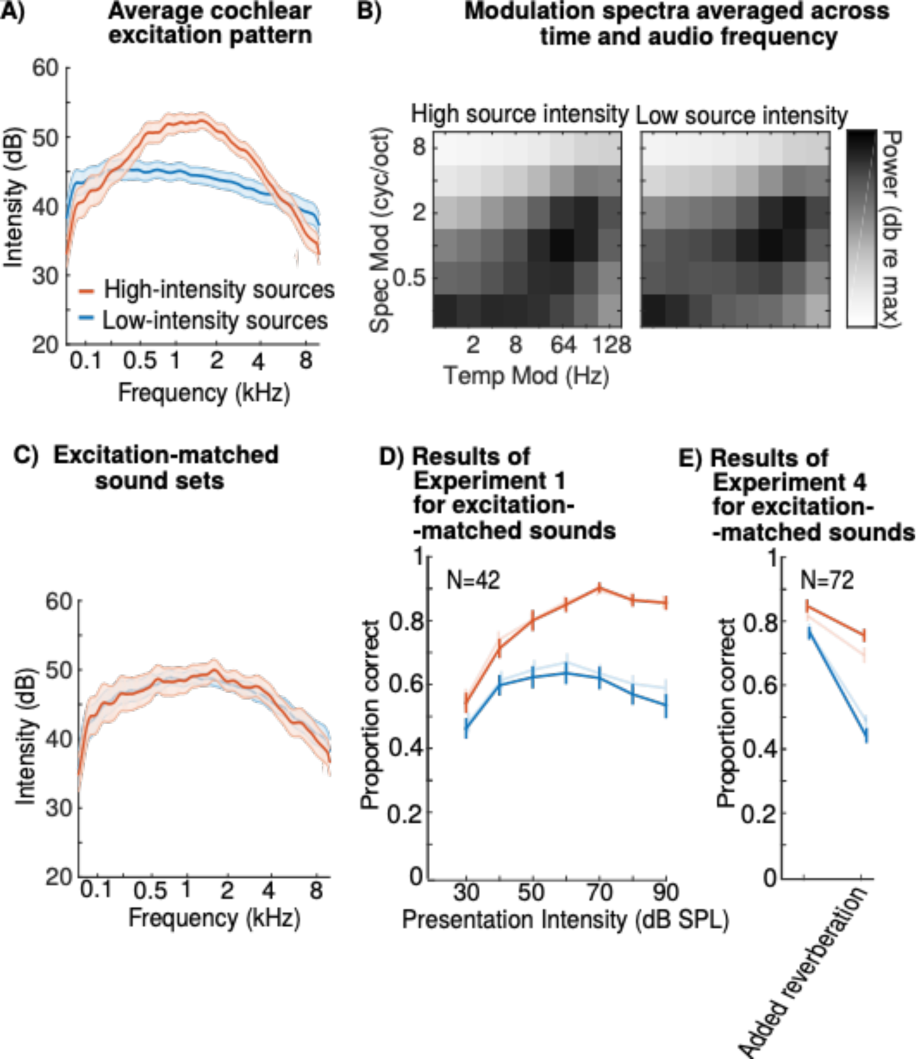
Source recognition effects from Figures 2 and 3 are not driven by differences in standard acoustic features. (A) The average spectral power distribution for low-intensity and high-intensity sources, measured using a Gammatone filter bank model of cochlear responses. We computed the envelope of each frequency band, converted the envelopes to a dB scale, averaged across time, and averaged across the sounds from each source intensity group. The graph plots the mean and standard deviation for each frequency band across the sounds from each group. (B) The strength of temporal and spectral modulations for sounds from each intensity group. Spectrotemporal modulations were computed by convolving a cochleagram with filters tuned to different rates of spectral/temporal modulation. Here we plot standard deviation of the filter responses over time, averaged across audio frequency. (C) To ensure that the differences in the spectral power distribution for low-intensity and high- intensity sources could not explain the differences in the effect of presentation intensity on their recognition, we selected a subset of 50 sounds from each group with closely matched excitation patterns. The graph plots the mean and standard deviation for each frequency band for these subsets. Note that the curves match well enough (as desired) that the blue curve is obscured by the red curve. (D&E) Results of Experiment 1 (recognition vs. presentation intensity) and Experiment 4 for subsets of low-intensity and high-intensity sources approximately equated for their spectral power distribution. The results are similar to those for the full sets of sounds (shown transparent), indicating that differences in the excitation patterns between the two sets of sounds do not account for the recognition differences.

We found that low-intensity and high-intensity sources had fairly similar modulation spectra, but that there were differences in the average excitation pattern, plausibly due to greater reverberation in the high-intensity sources, which tends to enhance mid-frequencies (Traer & McDermott, 2016). However, the interaction between presentation intensity and source intensity persisted for subsets of low- and high-intensity sources selected to yield matched average excitation patterns (Fig. 7C&D; F(6, 246) = 6.15; p < 0.001;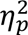 =0.130) (matching was performed by greedily discarding sounds so as to minimize differences in the excitation pattern). The interaction between reverberation and source intensity also persisted for the sound subsets matched in average excitation patterns (Fig 7E; F(1, 15) = 81.7; p < 0.001;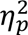=0.845). In addition, we ensured that our sound presentation system was linear over the range of intensities we presented, such that the results are unlikely to reflect distortion of sounds at high intensities. The results thus seem unlikely to reflect acoustic differences in the sounds tested.

## 16. Experiment 12: Sound recognition results are not driven by audibility

For low-intensity sources, many of the constituent frequencies may be inaudible in real-world listening conditions (Fig. 8A). If a recording of a low-intensity source is presented at a high intensity these frequencies may become audible and could potentially interfere with recognition by creating an unfamiliar acoustic profile. To test whether the unmasking of typically inaudible frequencies could explain our results, we used masking noise to prevent sound elements that were inaudible at low intensities from becoming audible at high intensities (Fig. 8A).

**Figure 8.**
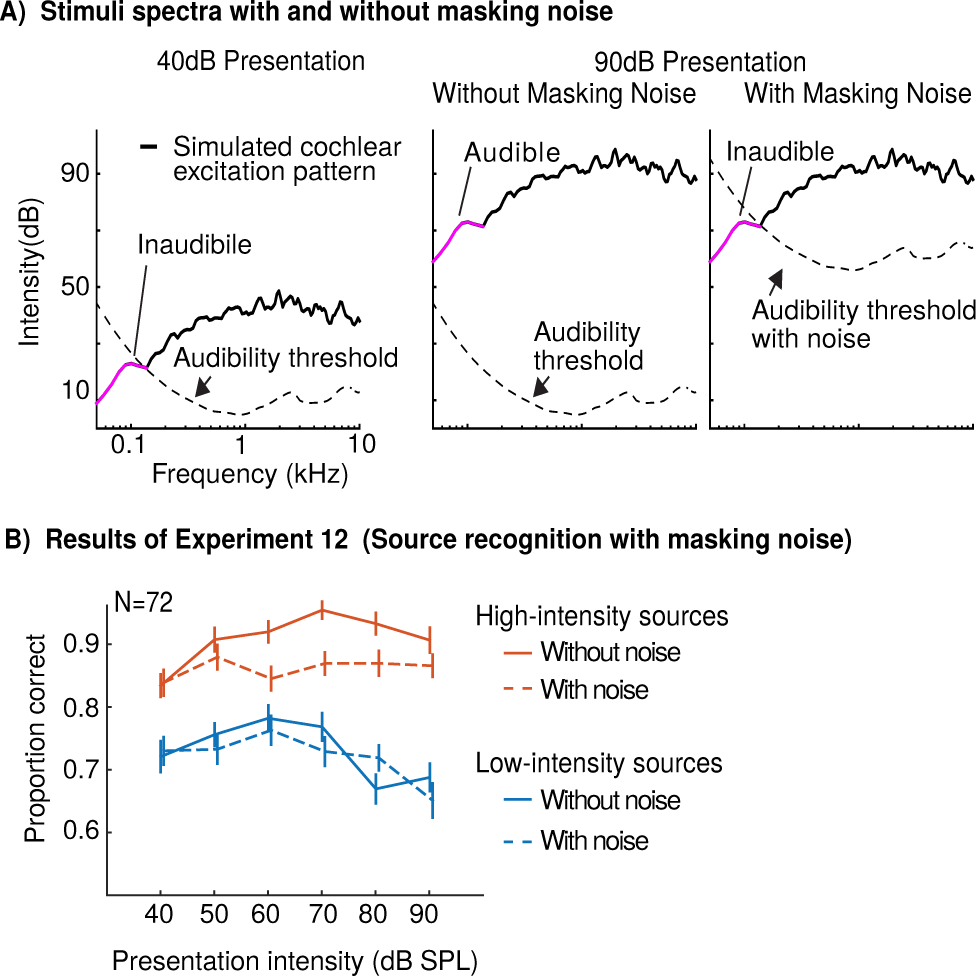
Audibility does not explain intensity-dependent recognition. (A) Illustration of the effect of overall sound intensity on audibility, and the use of masking noise to create stimuli with equal audibility profiles across different intensities. Each panel plots the maximum energy across time in each of a set of frequency channels (computed using a Gammatone filter bank model of cochlear responses) for a natural sound (“crumpling paper”) presented at two different overall intensities. Frequency-dependent audibility thresholds are plotted for comparison. At low intensities (left), some frequencies are below threshold. At high sound intensities (middle), these frequencies become audible. Such frequencies are presumably rarely heard for low-intensity sources, and could in principle interfere with their recognition when they become audible at high presentation intensities. Masking noise was used to keep these frequencies from becoming audible by elevating the threshold of audibility (right) (see Figure S2 and the Methods for a description of the masking noise). We note that the filtering of the cochlea is dependent on level, with bandwidths becoming somewhat broader with level (Glasberg and Moore, 2000), such that the excitation pattern at high levels is not simply a translated copy of the excitation pattern at low levels. However, we confirmed that the masking noise had the intended effect in a control experiment shown in Supplementary Figure S4, indicating that the assumption of a constant excitation pattern was sufficient to derive masking noise that had the intended effect. (B) The effect of masking noise on the recognition of low-intensity and high-intensity sources (same task as Experiment 1). Error bars show one standard error of the mean across subjects. The noise had no significant effect on the recognition of low-intensity sources, suggesting that audibility of normally inaudible frequencies was not the cause of their poor recognition at high presentation intensities. The noise impaired recognition of high-intensity sources, presumably due to the masking of frequencies that are often heard in daily life.

Experiment 12 was similar to Experiment 1 except that each sound was presented with and without masking noise. Sounds were presented at one of 6 intensities (40, 50, 60, 70, 80, and 90 dB SPL). Unlike in Experiment 1, we did not present sounds at 30 dB because performance was poor for both classes of sounds in this condition of Experiment 1. The masking noise was designed such that sound components that would normally be inaudible at 40 dB would remain inaudible at higher intensities (Fig. 8A). We confirmed that the masking noise had the intended effect in a supplementary experiment (described in the Appendix F).

### 16.1. Method

Experiment 4 was similar to Experiment 1 except that half of the trials included background noise designed to mask frequencies that were inaudible at the lowest presentation level. All other differences between the experiments are noted below.

#### 16.1.1 Participants

72 in-lab listeners participated in the experiment (44 female; mean age = 25.0, SD = 5.9) and had pure tone detection thresholds at or below 30 dB HL at all frequencies tested.

#### 16.1.2 Procedure

Each listener was presented with each of the 300 sounds once, at one of six presentation intensities (40, 50, 60, 70, 80, or 90 dB SPL) and either with or without masking noise. Across the 72 in-lab listeners, each sound was presented an equal number of times at each intensity (as in Experiment 1) and in each of the two noise conditions (with and without). The 21,600 descriptions provided by these 72 listeners (72 x 300 trials) were scored by 923 Mechanical Turk workers, each of whom scored 50 descriptions. As a consequence, most descriptions were scored by approximately two workers as in Experiment 1. The pattern of mean recognition performance across all conditions was again stable across independent sets of Mechanical Turk scorings (split-half Pearson correlation was 0.96).

#### 16.1.3 Masking Noise

The goal of the noise was to elevate the threshold of audibility such that frequencies that would be inaudible in quiet when sounds were presented at 40 dB (the lowest intensity condition in the experiment) would remain so at higher presentation intensities (see Fig. S2 for a schematic). We adapted threshold equalizing noise (TEN) (Moore, Huss, Vickers, Glasberg, & Alcántara, 2000), which equalizes the threshold of audibility for all frequencies (Fig. S2, middle panel). In our case, we wanted to elevate the threshold of audibility but maintain its dependence on frequency, and thus we shaped the spectrum of threshold-equalizing noise by the audibility threshold contour in quiet (Glasberg & Moore, 2006). Given that audibility varies smoothly with frequency on the scale of cochlear filter bandwidths, altering the noise spectrum in this way would be expected to cause detection thresholds to vary according to the audibility contour. The spectral shaping was accomplished in the frequency domain (using FFT/iFFT, interpolating the audibility contour to the grid of values sampled by the FFT, and multiplying the noise spectrum by the interpolated audibility contour). The overall level of the masking noise was set such that the resulting audibility threshold was (X-40) dB above the audibility threshold in quiet, where X is the overall intensity level of the stimulus. For the 40 dB condition, we would expect the noise to have little to no effect on audibility (although the noise itself was audible), and indeed the noise had no significant effect on performance in our discrimination task at this sound intensity (Fig. 5; Fig. S3; F(1, 71) = 0.09, p = 0.76). The effect of the masking noise was further validated in Experiment S1.

The noise had power between 50 Hz and 10 kHz (the Nyquist limit). We attenuated (by 60 dB) frequencies in the natural sounds that fell below the lower frequency cutoff of the noise. This attenuation was implemented in the frequency domain (using the FFT/iFFT), and we used a gradual roll-off rather than a sharp cutoff at 50 Hz to avoid unwanted time-domain effects (implemented by smoothing the ideal step filter with a Gaussian on a logarithmic frequency scale; FWHM=0.1 octaves).

#### 16.1.4 Stimulus Spectrum

In all other in-lab experiments (Experiments 1, 2, and 4), we used the audio transfer function of the headphones to adjust sound waveforms to have the desired overall level at the eardrum, but we did not otherwise compensate for the transfer function of the sound presentation system (i.e., so that the level of each frequency at the eardrum would correspond to its level in the recording). In Experiment 12 (and Experiment S1), we filtered the natural sounds and the masking noise in the frequency domain so that the power spectrum at the eardrum would match that of the original sound waveform. In practice, we found that the filtered natural sounds were perceptually very similar to the unfiltered natural sounds, and we observed similar effects of stimulus intensity in the absence of masking noise, suggesting that compensating for the system transfer function is not critical.

### 16.2 Results and discussion

In the absence of masking noise, we replicated our findings from Experiment 1 (Fig. 8B, solid lines). Recognition of high-intensity sources was good at moderate to high presentation intensities, while recognition of low-intensity sources declined at high presentation intensities, leading to an interaction between source intensity and presentation intensity (F(5, 355) = 4.70; p < 0.001; 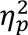 =0.062). If poor performance for low-intensity sources at high presentation intensities was due to unmasking of frequencies that are normally inaudible, then we would expect masking noise to eliminate this impairment. Alternatively, if the impairment reflects the inference of the source intensity, the masking noise should have little effect, as the overall sound intensity is what should matter. Under either account, it seemed plausible that the masking noise would impair performance for typically loud sources because the noise masks frequencies that, for high-intensity sources, are normally heard and could be used to aid recognition.

As shown in Fig. 8B (dashed lines), there was no significant effect of the masking noise for low-intensity sources (F(1, 71) = 0.33; p = 0.57; 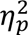=0.002), indicating that the recognition impairments we observe are not driven by audibility of normally inaudible sound components. By contrast, there was a small decrement in overall performance for high-intensity sources when masking noise was present (F(1, 71) = 20.00; p < 0.001; 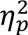=0.057; we observed intermediate results for sources with intermediate typical intensities; Fig. S3). As a consequence, the interaction between source intensity and presentation intensity remained even with the masking noise (F(5, 355) = 2.25; p < 0.05; 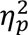=0.031). Thus, our findings suggest that the unmasking of inaudible frequencies cannot explain our results.

## 17. General discussion

A hallmark of human recognition is its robustness to the substantial acoustic variation created by different real-world environments. As a case study of how human listeners achieve robust recognition, we measured the extent to which recognition was invariant to sound intensity and reverberation, two variables that could plausibly be separated from the representation of a sound’s identity. A first set of experiments suggested that humans are not invariant to intensity (Experiments 1-3; Fig. 2). Sounds that do not normally occur at high intensities were often misidentified when presented at high intensities. This basic result replicated across several different experiments and could not be explained by simple acoustic features (Fig. 7) nor by changes in the audible frequency content of the sounds (Experiment 12; Fig. 8).

A second set of experiments revealed that reverberation implying a distant source had a similar effect on human recognition as high presentation intensity (Experiments 4-9; Figs. 3, 4 and 5). Low-intensity sources were misidentified in reverberation, while recognition of high-intensity sources was relatively robust. However, these failures of invariance were systematic: most errors were due to listeners mistakenly identifying low-intensity sources as high-intensity sources. By contrast, there was no evidence that sounds were misidentified more when presented with reverberation implying an atypical location for the source (i.e., when typically outdoor sources were convolved with reverberation typical of indoor spaces; Experiments 10&11, Fig. 6). Collectively the results indicate that sound recognition is not invariant to intensity or reverberation. Instead, the results are consistent with intuitive causal inference, in which the intensity of a sound source is implicitly inferred and used to constrain recognition judgments. Although not producing invariance across arbitrary stimulus manipulations of intensity of reverberation, this strategy likely aids accurate recognition in everyday settings, in which observed sounds must be physically consistent with their sources in the world.

We note that the experimental conditions were intended to maximize the chances of observing invariance. In experiments manipulating presentation intensity (i.e., Experiments 1, 2 and 12), listeners were told that sounds would be presented over a wide range of intensities, and in reverberation experiments (i.e., Experiments 4, 6-11), listeners were told that levels were artificially normalized, such that listeners should have been maximally inclined to benefit from any invariance mechanisms, and from decision strategies that accounted for the unrealistic presentation intensities. The fact that listeners were informed of the experimental design, and then experienced a wide range of intensities during the experiment makes it unlikely that listeners mistakenly assumed that sounds were played at veridical intensities, such that their mistakes reflect a counterproductive decision strategy. If anything, the effects we documented may have been weakened as a consequence of listeners’ knowledge of the experiment structure. One might also expect listeners to be somewhat adapted to variation in presentation intensities due to the common presence of electrical audio amplification in modern listening conditions, where playback levels of radios, televisions and other devices are independent of lawful physical relations related to a scene in the world. Such adaptation would also have weakened the effects described here. The differences we observed between low-intensity and high-intensity sources also do not appear to be due to low-level acoustic differences (Fig. 7), or to differences in recording conditions (Fig. 4D,E).

### 17.1 Prior work on invariance

Prior work suggests several reasons why listeners might be invariant to intensity. Gain control mechanisms exist as early as the cochlea (Darrow, Maison, & Liberman, 2006; Guinan, 2006) and midbrain (Dean, Harper, & McAlpine, 2005) that partially attenuate the effects of sound intensity. Moreover, mechanisms for level-invariant representation have been proposed at the level of the cortex in non-human animals (Billimoria et al., 2008; Sadagopan & Wang, 2008). In addition, many sounds occur over a wide range of levels in everyday experience (because we encounter them at a range of distances, or because the source can vary in physical intensity), such that a general normalization mechanism might be expected to emerge during auditory development. The fact that recognition is nonetheless influenced by cues to a source’s intensity is thus suggestive that inferred physical variables figure prominently in environmental sound recognition.

### 17.2 Limitations

The use of real-world sounds increases the relevance of our results for everyday hearing, but also presents methodological challenges (Shafiro, 2008; Shafiro & Gygi, 2004). Many real-world sound sources cannot practically be recorded in an anechoic environment (e.g., plane taking off, shower, truck, crowd cheering, gunshots, stream, traffic, rain, etc.), and are thus inevitably “contaminated” with reverberation and background noise. We dealt with this issue by using two sets of sounds: a large and diverse set of natural recordings, and a set of controlled studio recordings that was necessarily more limited in size and scope.

The uncontrolled reverberation in the natural set is likely to have affected listeners’ distance judgments (Experiment 5) and interfered with the use of added reverberation to manipulate distance. We were able to partially mitigate this by using distance-matched subsets of sources, and by replicating the results of some experiments using studio recordings. The similar effects evident across both sound sets suggest that our main results are robust to the reverberation originally present in the natural recordings (once equated for distance) and to the idiosyncrasies of the particular sounds in our studio recording set.

The studio recordings were limited by practical constraints. Many high-intensity and outdoor sounds were impossible to record, and many sounds we did record nonetheless entailed practical difficulties (e.g., chainsaw, lawnmower, chopping wood, walking on sand, shoveling, glass smashing). We were pleased to complete these recordings while avoiding damage to life or limb, or to our sound booth.

The studio recordings all had similarly minimal reverberation, but nonetheless produced variation in rated distance. This variation presumably reflects the influence of source knowledge, demonstrating another challenge of using recognizable sounds for which listeners have expectations and prior knowledge. We controlled for these effects again by using distance-matched subsets of sounds.

Another challenge is that we cannot obtain measurements of the intensities with which most real-world sounds are encountered in the world. In lieu of this we had humans rate whether sounds were typically soft or loud. These ratings are surely not perfectly reflective of actual source intensities. However, our analyses relied on very coarse divisions of sounds based on these ratings (e.g., into the loudest and quietest sources), and it seems likely that these divisions indeed capture substantial differences in source intensity. Moreover, we replicated our results using studio recordings where the true source intensities were known.

Similar issues were present for reverberation, where we asked humans to rate whether sounds are typically encountered indoors or outdoors in lieu of measuring reverberation. Typical outdoor environments have much less reverberation (i.e., shorter RT60s) than typical indoor spaces when source-listener distance is moderate (i.e., several meters) (Traer & McDermott, 2016). However, extremely high-intensity sources can be heard over kilometer-scales (e.g., thunder, gunshots, helicopters) and reverberation over such scales can have long decay times, possibly due to atmospheric turbulence as well as reflections (Knudsen, 1946). Although outdoor reverberation over such scales has not been characterized in detail, it is plausible that humans may encounter very distant, high-intensity sources in outdoor settings with reverberation that is similar to the “indoor” reverberation that we simulated. This may explain the typicality ratings we obtained in Experiments 10 and 11, in which added reverberation did not decrease the perceived appropriateness of the acoustic environment for high-intensity outdoor sounds (Fig 7B). Nonetheless, for lower-intensity sources, our synthetic reverberation produced a larger decrement in appropriateness for outdoor than indoor sources, presumably because low-intensity sources are typically encountered with such reverberation only when indoors. It would clearly be ideal to eventually substantiate this argument with measurements of reverberation from a large corpus of real-world audio and more thorough investigations of reverberation over large distances in outdoor scenes.

We note also that our manipulations of reverberation did not explore the full space of reverberation, instead using a single decay time and direct-to-reverberant ratio. We have no reason to think that the results are specific to the particular reverberation we used, but there is clearly room for a more exhaustive exploration of the effects of reverberation on recognition.

We relied exclusively on recognition accuracy and analyses of recognition confusions, but note that other experimental measures might give further insight. For instance, listening effort (Winn, Wendt, Koelewijn, & Kuchinsky, 2018) might also vary depending on whether sounds are presented in ecologically valid conditions, and might be more sensitive than accuracy. Inferred source properties might also be decodable from neurophysiological measurements.

### 17.3 Environmental sound perception

In general, human perception of environmental sounds has been little studied in comparison to speech or music, even though such sounds figure prominently in everyday behavior. Past studies have begun to characterize human environmental sound recognition, and have identified some of the acoustical features underlying this recognition (Balas, 1993; Gygi et al., 2004, 2007; McDermott, Schemitsch, & Simoncelli, 2013; McDermott & Simoncelli, 2011; McWalter & McDermott, 2018). In some cases listeners prefer to categorize environmental sounds according to their source than to acoustical features (Gygi et al., 2007; Lemaitre, Houix, Misdariis, & Susini, 2010). This latter finding is consistent with our hypothesis that recognition involves estimation of the properties of a sound source.

Our work also brings to light the importance of using ecologically valid intensities and reverberation in experiments with environmental sounds. By default one might be inclined to equate intensity (and/or reverberation) across experimental stimuli, but our results suggest this could have unintended consequences (e.g., if low-intensity sources are presented at moderate SPL levels, or with reverberation implying an implausibly large distance). We also highlight some of the challenges involved in using real-world sounds as stimuli. The distance judgments (Experiment 5, when compared with Experiment 7) suggest that real-world recordings likely carry reverberant cues to distance. Our experiments show that these cues indirectly affect the perceived source intensity. Source recognition in turn, may affect distance judgments (Fig 5C).

These implicit interactions merit consideration in experiment design when using natural sound recordings.

It is plausible that analogous effects exist for other properties that are associated with a source. For instance, some sounds tend to occur in particular locations relative to the listener (e.g., birds that mostly fly overhead (Parise, Knorre, & Ernst, 2014), or footsteps, which tend to come from below). Similarly, some sounds are much more likely to occur far from a listener than nearby. Such properties may also constrain recognition, and may also need to be considered when designing experiments.

### 17.4 Causal inference in audition

Our results suggest that listeners infer the underlying physical parameters that produce environmental sounds and use these parameters to recognize them. In the case of source intensity, the auditory system appears to jointly infer the distance and source intensity of a sound, and then infers a type of source consistent with this estimated source intensity. The role of distance cues has been previously noted in loudness judgments, which are proposed to reflect inferred source intensity (Zahorik & Wightman, 2001). However, the importance of such inferences for recognition, whether due to intensity or reverberation, had not been addressed prior to this paper. Our main contribution is to demonstrate that causal inferences have objectively measurable consequences on auditory recognition (arguably the most important auditory behavior), even for the simplest physical attributes of sound that one might naively think would be ignored for the purposes of recognition, particularly given the ubiquity of normalization processes in sensory systems (Carandini & Heeger, 2011; Schwartz & Simoncelli, 2001).

Examples of causal inference (i.e., estimation of a causal parameter in a non-trivial generative model) are fairly well established in various aspects of vision: object recognition (Kersten, Mamassian, & Yuille, 2004), shape from shading (Adams, Graf, & Ernst, 2004), size estimation (Oyama, 1974), audiovisual integration (Shams & Beierholm, 2010), and intuitive physical judgments (Gerstenberg, Goodman, Lagnado, & Tenenbaum, 2012). Our work suggests that sophisticated implicit inferences are also fundamental to human audition.

## Acknowledgements

The authors thank Dekel Zeldov and Selena Wang for assistance running experiments, Jeremy Schwartz for assistance recording sounds, and Leyla Isik, Jackson Graves, and Ratan Murty for comments on an early version of the manuscript. This work was supported by a McDonnell Scholar Award to JHM and NSF Grant No. BCS-1454094, NSF Grant No. BCS- 1921501, NSF Grant No. PHY17-48958 and NIH Grant No. R25GM067110. SNH was funded by an NSF graduate research fellowship and a postdoctoral fellowship from the Howard Hughes Medical Institute through the Life Sciences Research Foundation.

## Supplementary material

Sound stimuli and responses from listeners and online graders for all experiments are available at https://github.com/jt-uiowa/causal-inference-in-environmental-sound-recognition_data

## Appendix A. Source Intensity and Location Ratings

### A.1 Intensity

As a proxy for the typical physical source intensity for each real-world sound in our set (which would be impractical to measure), we asked a second set of online workers to rate the typical intensity of a sound on a scale of 1-10 using the following instructions:

*Listen to each of the sounds presented below and indicate how loudly you typically hear each sound in your daily life on a 1 (quiet) to 10 (loud) scale. For example, sounds typically heard at very quiet sound level such as writing (pen on paper) or typing would be rated as a 1, while sounds that are typically very loud, such as a jet engine or a jack-hammer would be rated as a 10. For sounds that can be heard at a variety of sound levels, indicate the level at which you most frequently hear that sound*.

We collected ratings from 389 online workers, which was sufficient to produce split-half correlations of the average rating for a sound that exceeded 0.9.

We chose to present the online workers with both the sound and a description of the sound so that they would have a good sense for the type of sound about which were asking. In this experiment, all sounds were normalized to have the same root-mean-square amplitude. The absolute intensity was set individually by workers because they listened to sounds on their own devices.

These ratings were averaged across online workers and used to classify sound clips into low-intensity and high-intensity sources, on the assumption that sounds that are typically loud in everyday life tend to be produced by high-intensity sources, whereas those that are typically quiet tend to be produced by low-intensity sources.

### A.2 Location

The sounds were also divided into two groups of typically indoor and typically outdoor sounds. To do this we asked a set of online workers to listen to each of the 300 recorded sounds and rate (on a scale of 1-10) how likely the sound was to be encountered indoors as opposed to outdoors. Workers received the following instructions:

*Listen to each of the sounds presented below and indicate how likely you are to encounter this sound in an indoor environment as opposed to an outdoor environment. Use a 1 (only heard outdoor) to 10 (only hear indoor) scale. For sounds that can be both indoors or outdoors, indicate how likely you think the sound is to be heard indoors*.

We collected ratings from 50 online workers. We presented the online workers with the sound recording as well as a description of the sound. The sounds were categorized by dividing them into two groups (typically indoor and typically outdoor sounds) around the median value.

## Appendix B. Headphone Calibration

For in-lab listening experiments (Experiments 1, 2, 4 and 12), sounds were presented through Sennheiser HD280 Pro headphones, which we calibrated using a Svantek 979 sound meter attached to a GRAS microphone with an ear and cheek simulator (Type 43-AG). We used this setup to estimate the transfer function of our entire sound presentation system (from the computer to the eardrum), by playing pink Gaussian noise and comparing the input spectrum with the spectrum measured by the microphone. We used this frequency response to calculate the overall sound pressure level of a sound for a given input waveform (by computing the power spectrum of the original waveform, multiplying by the gain at each frequency, and then summing the adjusted power across frequencies), and then scaled the waveform so as to yield the desired sound pressure level at the ear.

All other experiments were conducted entirely online because they either had no listening component (Experiments 3, 5, and 9), or because they were very short (Experiments 6-8 and 10-11), which made in-lab recruitment difficult.

## Appendix C. Hearing Thresholds

A majority (56%) of listeners in Experiment 1 had pure tone detection thresholds at or below 30 dB HL at all frequencies tested (0.25 to 8 kHz), but some of the older listeners had elevated thresholds, typically at higher frequencies. In Experiment 12, we tested a younger cohort all of whom had hearing thresholds below 30 dB to ensure that the results were robust to incidental hearing impairment.

In other in-lab experiments where all sounds were presented at 70dB, and in all online listening experiments, we did not measure detection thresholds as sounds were intended to be well above threshold. All listeners in these experiments self-reported normal hearing. All online listening tasks included a test to ensure that listeners were wearing headphones (Woods et al., 2017).

## Appendix D. Studio Recordings

Forty-five sound sources were recorded in a soundproof booth that was heavily-damped to minimize reverberation (Table 2). All recordings were made with a microphone positioned 10cm from the sound source (a Rode NT1A with a Focusrite Scarlett 6i6 Analogue-to-Digital-Converter). SPL measurements were made for each source with a Svantek SVAN 979 Sound & Vibration Analyzer (also positioned with a 10cm source-recorder separation). The sources were chosen to span a range of SPL levels and typical source locations. The gain settings were adjusted for each recording to capture an appropriate dynamic range for the sound. From several minutes of recordings for each sound, 5-second snippets were extracted. Where possible the snippet was chosen without truncating the sound (e.g., for the hatchet striking a log a snippet might contain one or two impacts). For continuous sounds (e.g., lawnmower) the snippet was given 200ms linear fades in and out. In each experiment one of these snippets was randomly selected for each source. For each of the snippets of audio obtained, several hours were spent preparing, cleaning, and ventilating the soundproof booths.

In addition to the sounds shown in Table 2, five additional sounds were recorded that were intermediate in both intensity and location (Sawing wood (81dB); Ratchet wrench (75dB); Drawing a nail from a box (nails scraping and sliding) (69dB); Velcro (66dB); Coin dropped on hard surface (65dB)). These sounds were presented in experiments but omitted from analysis in the interest of brevity and clarity.

## Appendix E. Acoustic Analyses

We assessed the extent to which low-intensity and high-intensity sources differ on standard acoustic features (Fig. 7). We first measured the average simulated cochlear “excitation pattern” for the different source-intensity groups, which is the average power across a filter bank that simulates cochlear filtering. Each sound waveform was convolved with a Gammatone filter bank (Slaney, 1998) (128 filters, with center frequencies between 20 and 10,000 Hz). We then computed the envelopes of the filter responses over time (via the Hilbert transform), converted these envelopes to a dB scale, and averaged these values across time and across sounds from the same source intensity group. We found that the differences between groups in their excitation pattern were modest relative to the variation within a group (Fig. 7A; error bars plot the standard deviation of the energy in each frequency bin across the sounds from a given intensity group).

To ensure that the modest differences in the mean excitation patterns of the sound groups could not explain the differences in their recognition, we analyzed recognition performance for subsets of the sounds that were approximately equated in their excitation patterns. We selected a subset of 50 low-intensity and 50 high-intensity sources with approximately matched excitation patterns by greedily discarding sounds from each group (starting from an initial pool of the full set of 75 sounds per group). At each iteration, we discarded the sound that led to the biggest reduction in the mean squared error between the average excitation patterns for the low-intensity and high-intensity source groups. We ran this algorithm for 50 iterations, alternating between discarding low-intensity and high-intensity sources, so as to discard 25 sounds per group. This was sufficient to produce similar average excitation patterns for the two groups (Fig. 7C).

We next measured the amount of temporal and spectral amplitude modulation, using a standard set of spectrotemporal modulation filters (Chi et al., 2005). Modulation was measured in cochleagrams computed using a filter bank similar to the Gammatone filter bank described above (116 filters between 50 and 10,000 Hz, with frequency responses shaped like the positive portion of a cosine function, with 87.5% overlap between adjacent filters; we used this filter bank for convenience because the modulation model described below was implemented using these filters). The envelopes of the cochlear filter responses were compressed to capture the effects of cochlear amplification at low intensity levels (by raising them to the power of 0.3). The resulting cochleagram was then convolved in time and frequency with spectrotemporal filters tuned to each of 9 different temporal modulation rates (0.5 to 128 Hz in octave steps) and 6 different spectral modulation scales (0.25 to 8 cycles per octave in octave steps). All of the filters were bandpass, and their properties have been described previously (Chi et al., 2005). The output of the modulation filter bank was a 4D tensor measuring energy in the sound as a function of time, audio frequency, temporal modulation rate and spectral modulation scale. We computed the standard deviation across time of this tensor (as a measure of the strength of the temporal fluctuations in each filter’s response), averaged across audio frequency, and averaged across sounds from a given intensity group. The result is a 2D matrix which represents the average energy of fluctuations at different temporal and spectral modulation rates (Fig. 7B). We found that the pattern of temporal and spectral modulations was similar between the different real-world intensity groups.

## Appendix F. Experiment S1: Verifying the Masking Effects of the Noise from Experiment 4

The masking noise in Experiment 12 was designed using pure tone audibility thresholds (Moore et al., 2000), and thus it was not obvious a priori that it would have the desired effect when used with natural sounds. We therefore performed a control experiment to verify that the noise had the desired effect. This experiment is somewhat complex, and because the results confirmed that the noise had the desired effect on audibility, the casual reader need not feel obligated to labor through the details.

We tested the effectiveness of the masking noise using a discrimination paradigm in which we attenuated low-intensity frequencies from natural sounds (details below) and assessed whether listeners could detect their absence with and without masking noise (Fig. S4A). On each trial listeners were asked to judge which of two intervals contained different sounds: in one interval, the same unaltered natural sound was presented twice, and in the other interval, the unaltered version was followed by a filtered version with low-intensity frequencies attenuated (by 30 dB). We expected that the change to the spectrum would be most noticeable at higher overall sound intensities, where more of the spectrum would be audible (Fig. S4B).

The goal of Experiment S1 was to test whether this anticipated improvement at higher sound intensities would be eliminated by the use of masking noise designed to prevent additional frequencies from becoming audible at high intensities.

We presented sounds at three intensities (40, 75, 90 dB) with and without noise. For the lowest-intensity condition (40 dB), we attenuated frequencies based on their maximum power over time (computed from a cochleagram, described below) relative to the threshold of audibility (Fig. S4B). We used the maximum power over time (rather than, for example, the mean) because in principle listeners might detect energy in a frequency band any time it exceeds the audibility threshold. For the higher-intensity conditions (75 & 90 dB), we instead attenuated frequencies based on their maximum power relative to the elevated audibility threshold we intended to produce with noise. If the noise had the intended effect then it should have reduced performance on the high-intensity conditions to that of the 40 dB condition.

We measured the time-varying power of different frequency bands using a Gammatone filter bank designed to mimic the frequency tuning in the cochlea. We then attenuated (by 30 dB) all frequency channels whose maximum power over time fell below a certain “audibility-relative” cutoff (see Fig. S4C for an illustration). We manipulated difficulty by varying the cutoff, with higher cutoffs causing more of the spectrum to be suppressed and thus making the task easier. This approach allowed us to measure detection accuracy as a function of the cutoff for each condition in the experiment (Fig. S4D).

### F.1 Participants

Twenty-two listeners participated in the experiment (12 female; mean age = 25.4 years, SD = 3.3 years). All but one listener had pure tone detection thresholds at or below 30 dB HL. One listener had a threshold of 40 dB HL in their left ear at 3 and 4 kHz; the exclusion/inclusion of their data did not affect the results.

### F.2 Stimuli and Procedure

On each trial, listeners heard four presentations of a natural sound, divided into two intervals (Fig. 6A). In one interval, one of the two sounds was filtered to attenuate frequencies below an “audibility-relative” cutoff (as described in the results; details of the filtering are described below). Listeners were instructed to indicate the interval in which the two sounds differed. Each sound was 2 seconds in duration. There was a 600 ms gap in between sounds from the same interval, and a 1-second gap between the two intervals. Linear ramps (100 ms in duration) were applied to the beginning and end of each sound.

Sounds were presented at one of three intensities (40, 75, and 90 dB), with or without background masking noise, and with 5 different audibility-relative cutoffs (0, 5, 15, 25, and 35 dB). The masking noise was the same as that used in Experiment 4. The noise lasted throughout the duration of each trial (starting 500 ms before the first sound of the first interval and ending 500 ms after the offset of the last sound of the second interval). Each listener heard a different subset of 12 of the 300 natural sounds from Experiments 1 and 4. This relatively small number of sounds was chosen so that each sound could be presented once in each of the 30 different conditions (3 intensities x 5 cutoffs x 2 noise conditions – with and without), yielding 360 trials. We excluded sounds in which the power at most frequencies fell below the maximum audibility relative cutoff (35 dB; since this would have caused nearly the entire spectrum to be suppressed), leaving a pool of 186 sounds. The set of 12 sounds used for a listener was randomly drawn from this set of 186.

The experiment was divided into 12 sections of 30 trials, and after each section the listener was given the option to take a short break.

### F.3 Filtering

We used a Gammatone filter bank to model cochlear responses as a function of time and frequency (128 filters with center frequencies between 20 Hz and 22,050 Hz) (40). Sound waveforms were sampled at 44,100 Hz. We measured the Hilbert envelope of each filter’s output, and converted this envelope to dB SPL. For each filter, we computed whether its envelope for a given sound/condition fell above or below the audibility-relative cutoff, yielding a binary vector of zeros and ones indicating which frequencies to attenuate. To avoid time-domain artifacts (e.g., ringing), we smoothed this binary vector using a Gaussian kernel on a logarithmic frequency scale (FWHM = 0.1 octaves), yielding a new vector with smoothed values between 0 and 1. This vector was multiplied by 30. We then attenuated each frequency by the number of decibels specified in the corresponding element of the vector. The frequency attenuation was implemented in the frequency domain (using the FFT/iFFT and interpolating the attenuation vector to the frequencies sampled by the FFT).

### F.4 Results

Fig. S4D shows performance for the three most diagnostic conditions: sounds presented at 40 dB without noise, at 90 dB without noise, and at 90 dB with noise. As expected, performance increased as the cutoff was increased (F(4, 84) = 88.47, p < 0.01), and, in the absence of masking noise, performance increased with stimulus intensity (F(2, 42) = 37.61, p < 0.01). However, performance at 90 dB with noise was very similar to performance at 40 dB without noise. As a consequence, there was no significant effect of intensity on performance with noise (F(2, 42) = 1.63; p = 0.21), and there was a significant interaction between the effect of intensity and the effect of masking noise (F(2, 42) = 11.92, p < 0.01). These results suggest that most of the frequencies that became audible at 90 dB were masked by our noise, as intended.

## Supplementary Information

**Table S1.**
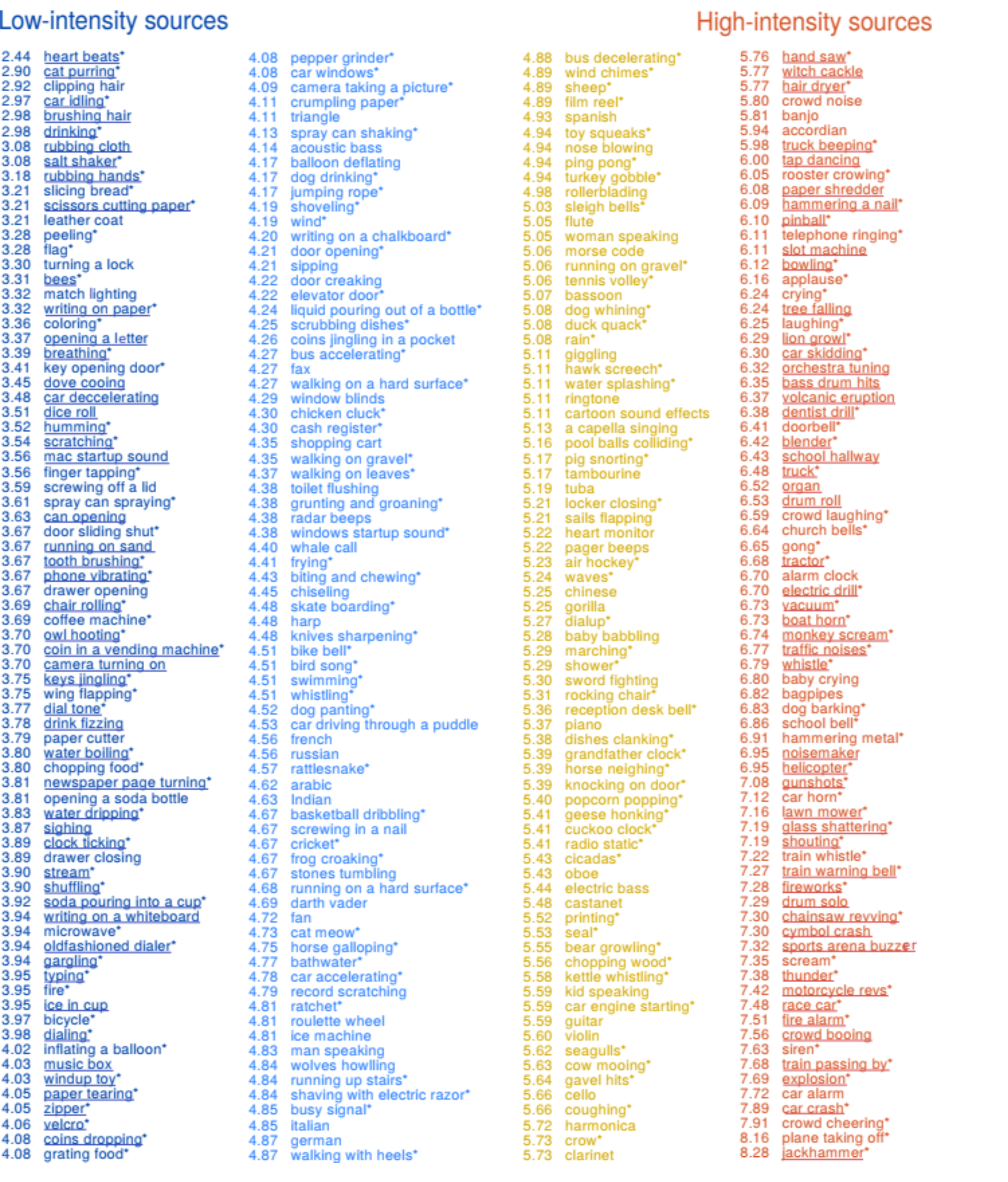
This table lists all 300 sounds tested in the experiments. Sounds were partitioned into 4 groups based on their rated real-world source intensity on a 1 (lowest-intensity) to 10 (highest) scale. The rated intensity is shown next to each sound. The color indicates the group the sound was assigned to. In Experiments 1-3 and 12 the left column of 75 sounds were classed as “low-intensity” and the right column of 75 sounds as “high-intensity”. The subset of sounds used in Experiments 4-5 are marked with asterisks (*). The difficulty matched-subsets (Fig 2C) are underlined. The distance-matched subsets used in Experiments 7-11 are listed in Tables S2-S4.

**Figure S1.**
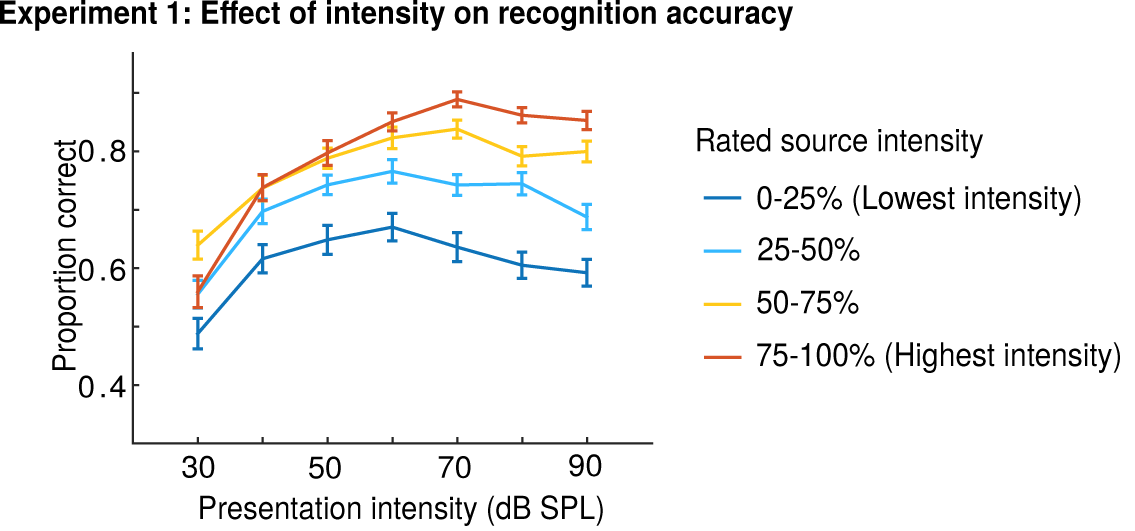
Same as Figure 2A but showing performance for all 4 partitions of the sound set based on their rated real-world source intensity.

**Figure S2.**
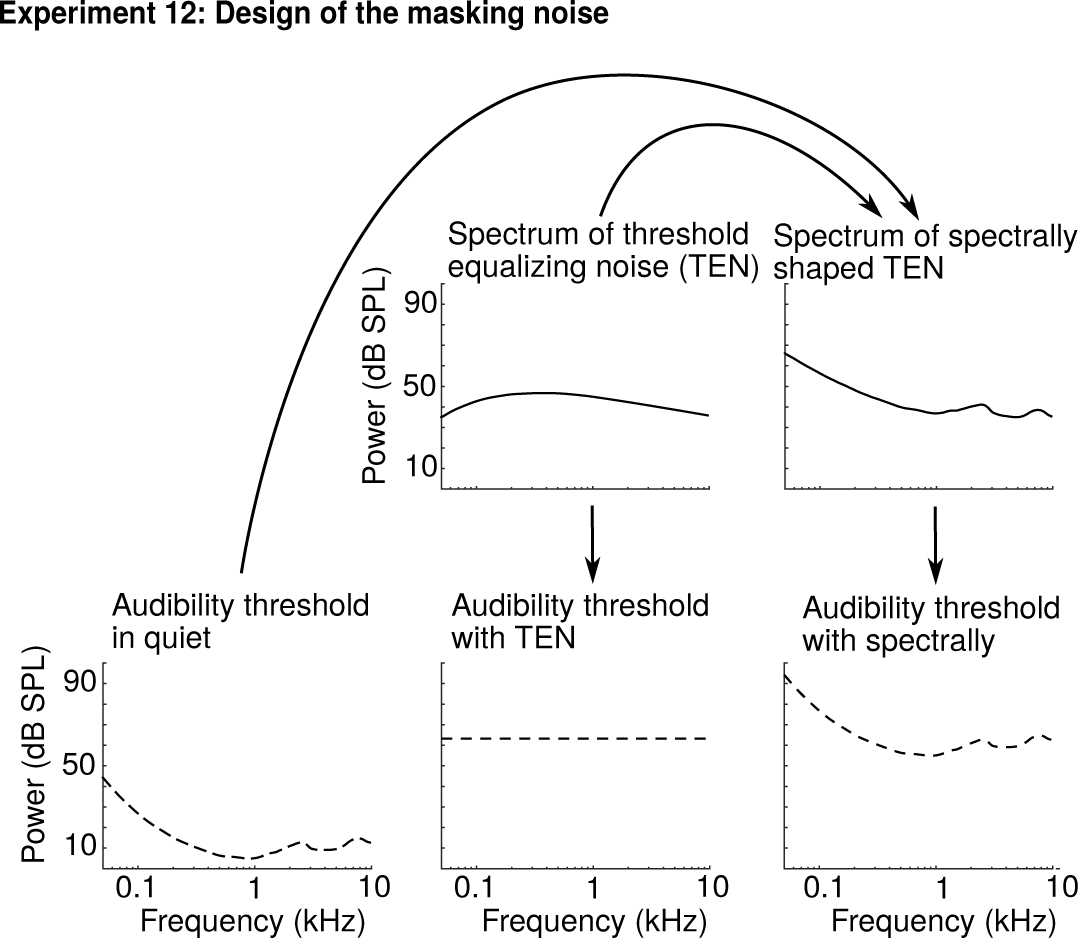
The goal of the masking noise was to elevate the audibility threshold so that frequencies that would normally be inaudible at low sound intensities remain inaudible at higher intensities. This goal was accomplished by starting with threshold equalizing noise (TEN) (41), which equates thresholds for all frequencies. We then shaped TEN with the contour of audibility in quiet so that the audibility threshold would be elevated rather than flattened. This figure plots the power spectrum (computed with the FFT) and expected audibility threshold for TEN and our spectrally shaped noise. In the experiment, the overall intensity of the masking noise was yoked to the intensity of the stimuli, causing the audibility threshold to shift up and down with the intensity of the stimulus. In this figure we show the spectrum and audibility curves corresponding to a single high-intensity stimulus (90 dB).

**Figure S3.**
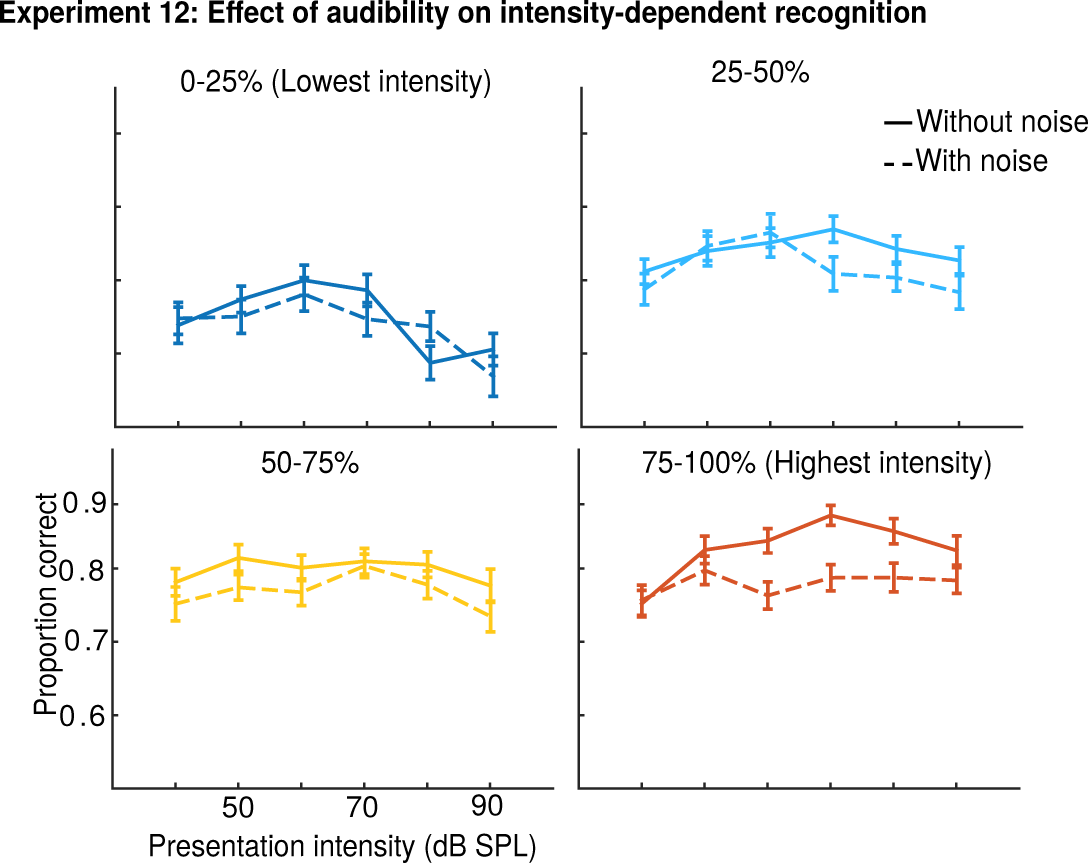
Same as Figure 8B but showing performance for all 4 partitions of the sound set based on their rated real-world source intensity.

**Figure S4.**
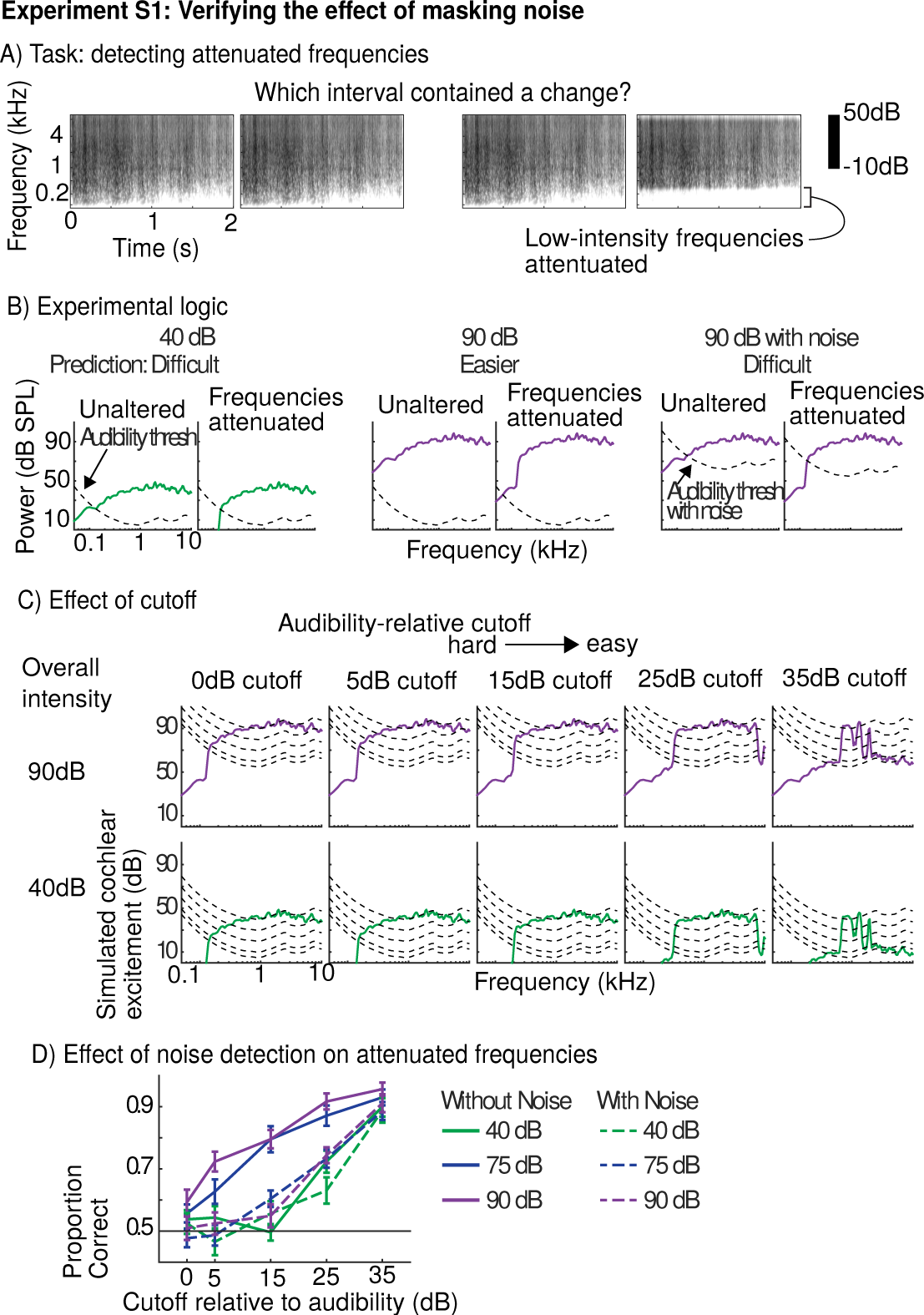
Design and results of Experiment S1: Validation of masking noise. (A) Schematic of the task used to assess the efficacy of the masking noise. Each trial comprised two intervals. In one interval the same natural sound was presented twice. In the other interval, one of the sounds was filtered to attenuate low-intensity frequencies. Listeners were asked to detect which interval contained a change between the two sounds. (B) The experiment was designed such that suppressed frequencies should be easier to detect for higher-intensity stimuli due to greater audibility. The masking noise was designed to eliminate this benefit by raising the audibility threshold. (C) Frequencies were attenuated that fell below a certain intensity cutoff relative to the threshold of audibility (see text for details). Higher cutoffs cause more frequencies to be attenuated, making the task easier. (D) Discrimination performance as a function of the cutoff for stimuli presented at 40 dB without noise, at 90 dB without noise, and at 90 dB with noise. Error bars show one standard error of the mean across subjects. As predicted, performance was substantially better for higher-intensity stimuli without noise, but this benefit was eliminated by the masking noise, demonstrating that the masking noise had the intended effect.

**Table S2:** Distance-matched groupings of low- and high-intensity sounds obtained from Experiment 6 and used in Figure 4. The numbers show the intensity ratings as given in Table S1.

| **<u>Low-intensity sounds</u>** | **<u>High-intensity sounds</u>** |
| --- | --- |
| car idling 2.97 | bus decelerating 4.88 |
| slicing bread 3.21 | sheep 4.89 |
| finger tapping 3.56 | Ping pong 4.94 |
| chair rolling 3.69 | turkey gobble 4.94 |
| coffee machine 3.69 | rain 5.08 |
| owl hooting 3.70 | water splashing 5.11 |
| dial tone 3.77 | pool balls colliding 5.16 |
| stream 3.90 | pig snorting 5.17 |
| gargling 3.94 | marching 5.29 |
| fire 3.95 | shower 5.29 |
| typing 3.95 | rocking chair 5.31 |
| grating food 4.08 | reception desk bell 5.36 |
| jumping rope 4.17 | dishes clanking 5.38 |
| shoveling 4.19 | cuckoo clock 5.41 |
| wind 4.19 | printing 5.52 |
| elevator door 4.22 | kettle whistling 5.58 |
| walking on gravel 4.35 | cow mooing 5.63 |
| walking on leaves 4.37 | coughing 5.66 |
| grunting and groaning 4.38 | hair dryer 5.77 |
| bike bell 4.51 | crying 6.24 |
| swimming 4.51 | laughing 6.25 |
| rattlesnake 4.57 | dentist drill 6.38 |
| frog croaking 4.67 | doorbell 6.41 |
| basketball dribbling 4.67 | blender 6.42 |
| horse galloping 4.75 | vacuum 6.73 |
| car accelerating 4.78 | dog barking 6.83 |

**Table S3:** Distance-matched groupings of low- and high-intensity studio-recorded sounds obtained from Experiment 7 and used in Figure 5.

| **<u>Low-intensity sounds</u>** | **<u>High-intensity sounds</u>** |
| --- | --- |
| <u>rustling branch (52dB)</u> | <u>electric can opener (83dB)</u> |
| <u>suitcase rolling (52dB)</u> | <u>coffee bean grinder (88dB)</u> |
| <u>branch trimmer (53dB)</u> | <u>bicycle bell (92dB)</u> |
| <u>footsteps in sand (55dB)</u> | <u>hair dryer (92dB)</u> |
| <u>pepper grinder (56dB)</u> | <u>dropping stones on stones (92dB)</u> |
| <u>biting into an apple (57dB)</u> | <u>compressed air spray 95dB</u> |
| <u>splashing water (57dB)</u> | <u>vacuum cleaner (95dB)</u> |
| <u>zipper (58dB)</u> | <u>hammering a nail into wood (97dB)</u> |
| <u>shoveling sand (59dB)</u> | <u>drill (101dB)</u> |
| <u>pouring liquid (59dB)</u> | <u>glass smashing (102dB)</u> |
| <u>peeling vegetables (59dB)</u> | <u>hammering metal (112dB)</u> |
| <u>chopping vegetables (60dB)</u> | <u>chainsaw (113dB)</u> |
| <u>scissors (62dB)</u> | <u>leaf blower (114dB)</u> |

**Table S4:** Distance-matched groupings of indoor and outdoor sounds used in Fig 6. The numbers show the intensity ratings as given in Table S1. “High-intensity” sounds are marked with an asterisk.

| **<u>Indoor sounds</u>** | **<u>Outdoor sounds</u>** |
| --- | --- |
| slicing bread 3.21 | breathing 3.39 |
| finger tapping 3.56 | spray can shaking 3.61 |
| coffee machine 3.69 | stream 3.90 |
| chair rolling 3.69 | fire 3.95 |
| dial tone 3.77 | jumping rope 4.17 |
| gargling 3.94 | shoveling 4.19 |
| grating food 4.08 | walking on gravel 4.35 |
| elevator door 4.22 | walking on leaves 4.37 |
| grunting and groaning 4.38 | bike bell 4.51 |
| running up stairs 4.84 | dog panting 4.52 |
| ping pong* 4.94 | rattlesnake 4.57 |
| shower* 5.29 | frog croaking 4.67 |
| dishes clanking* 5.38 | ratchet 4.81 |
| cuckoo clock* 5.41 | walking with heels 4.87 |
| coughing* 5.66 | bus decelerating* 4.88 |
| hair dryer* 5.77 | sheep* 4.89 |
| applause 6.16 | rain* 5.08 |
| crying* 6.24 | water splashing* 5.11 |
| laughing* 6.25 | pig snorting* 5.17 |
| dentist drill* 6.38 | marching* 5.29 |
| doorbell* 6.41 | horse neighing* 5.39 |
| blender* 6.42 | cicadas* 5.43 |
| vacuum* 6.73 | cow mooing* 5.63 |
| crowd laughing* 6.59 | crow* 5.73 |
| glass shattering* 7.19 | whistle* 6.79 |
| fire alarm* 7.51 | train warning bell* 7.27 |

**Table S5:** Distance-matched groupings of indoor and outdoor sounds obtained from Experiment 7 and used in Fig 6. “High-intensity” sounds are marked with an asterisk.

| **Indoor sounds** | **Outdoor sounds** |
| --- | --- |
| peeling vegetables (59dB) | rustling branch (52dB) |
| chopping vegetables (60dB) | branch trimmer (53dB) |
| scissors (62dB) | footsteps in sand (55dB) |
| crumpling paper (69dB) | splashing water (57dB) |
| electric shaver (72dB) | shoveling sand (59dB) |
| stapler* (75dB) | footsteps in pebbles (69dB) |
| clanking dishes* (78dB) | walking in dry leaves (69dB) |
| electric can opener* (83dB) | spray can shaking* (76dB) |
| hair dryer* (92dB) | hatchet striking a log* (78 dB) |
| vacuum* (95dB) | leaf blower* (114dB) |

